# MSH2-MSH3 promotes DNA end resection during HR and blocks TMEJ through interaction with SMARCAD1 and EXO1

**DOI:** 10.1101/2021.04.23.441074

**Authors:** Jung-Min Oh, Yujin Kang, Jumi Park, Yubin Sung, Dayoung Kim, Yuri Seo, Eun A Lee, Jae Sun Ra, Enkhzul Amarsanaa, Young-Un Park, Hongtae Kim, Orlando Schärer, Seung Woo Cho, Changwook Lee, Kei-ichi Takata, Ja Yil Lee, Kyungjae Myung

## Abstract

DNA double strand break (DSB) repair by Homologous recombination (HR) is initiated by the end resection, a process during which 3’ ssDNA overhangs are generated by the nucleolytic degradation. The extent of DNA end resection determines the choice of the DSB repair pathway. The role of several proteins including nucleases for end resection has been studied in detail. However, it is still unclear how the initial, nicked DNA generated by MRE11-RAD50-NBS1 is recognized and how subsequent proteins including EXO1 are recruited to DSB sites to facilitate extensive end resection. We found that the MutSβ (MSH2-MSH3) mismatch repair (MMR) complex is recruited to DSB sites by recognizing the initial nicked DNA at DSB sites through the interaction with the chromatin remodeling protein SMARCAD1. MSH2-MSH3 at DSB sites helps to recruit EXO1 for long-range resection and enhances its enzymatic activity. MSH2-MSH3 furthermore inhibits the access of DNA polymerase θ (POLQ), which promotes polymerase theta-mediated end-joining (TMEJ) of DSB. Collectively, our data show a direct role for MSH2-MSH3 in the initial stages of DSB repair by promoting end resection and influencing DSB repair pathway by favoring HR over TMEJ. Our findings extend the importance of MMR in DSB repair beyond established role in rejecting the invasion of sequences not perfectly homologous to template DNA during late-stage HR.

## Introduction

Genome integrity is constantly challenged by DNA replication errors and by diverse endogenous and exogenous damaging agents such as oxidative stress and environmental radiation (Friedberg et al., 2006; Jackson and Bartek, 2009; Rouse and Jackson, 2002; Su, 2006). To maintain genomic stability, various DNA repair mechanisms have evolved in cells. DNA mismatch repair (MMR) is a repair modality conserved in all organisms used to correct DNA mismatches and insertion-deletion loops (IDL) resulting from DNA replication errors and recombination between closely related, but not identical DNA sequences (Kolodner and Alani, 1994; Li, 2008). MMR is initiated by two types of MMR protein complexes, depending on the nature of DNA mismatches. In eukaryotes, MutSα is a heterodimer composed of MSH2 and MSH6 that recognizes mismatches of 1 or 2 nucleotides. MutSβ is composed of MSH2 and MSH3 spanning more than 2 nucleotides and IDLs (Drummond et al., 1995; Genschel et al., 1998; Habraken et al., 1996). Following MutS complexes binding to DNA lesions, MutL heterodimers (MLH1-PMS2, MLH1-PMS1 or MLH1-MLH3) are recruited with MutS to enhance mismatch recognition, and to promote a conformational change in MutS allowing for the MutL/MutS complex to slide away from mismatched DNA (Allen et al., 1997; Gradia et al., 1999). Repair is then initiated by a single-stranded nick generated by MutL at some distance to the lesion (Kadyrov et al., 2006; Kadyrov et al., 2007). Exonucleolytic activities involving exonuclease 1 (EXO1) remove the mismatch containing strand, and the correct genetic information is restored by gap filling (Fishel, 1999; Jiricny, 2006). Germline mutations of MMR genes cause Lynch syndrome, a hereditary disease primarily characterized by non-polyposis colorectal cancer. While Lynch syndrome patients are rare, approximately 20% of all colorectal cancers are MMR defective (Bronner et al., 1994; Fishel et al., 1993; Miyaki et al., 1997; Nicolaides et al., 1994; Papadopoulos et al., 1994). Knowing the MMR status of a cancer provides a chance for tailored chemotherapy, and for cancer immunotherapies taking advantage of the excessive number of neoantigens arising in MMR defective tumors (Devaud and Gallinger, 2013; Le et al., 2015; van Geel et al., 2015).

DNA double-strand breaks (DSB) are considered to be the most threatening form of DNA damage. Since unrepaired DSBs can cause chromosome discontinuity and translocations, proper and precise DSB repair is essential to prevent genomic instability, and ultimately tumorigenesis (Hanahan and Weinberg, 2011; Jackson and Bartek, 2009; Smeenk and van Attikum, 2013). DSBs are repaired by three major pathways; homologous recombination (HR), non-homologous end-joining (NHEJ) and microhomology-mediated end joining [MMEJ; largely conferred by DNA polymerase θ (POLQ) mediated end joining (TMEJ)] (Chang et al., 2017). A key factor determining the pathway choice between using HR, NHEJ or TMEJ pathways is the lenghth of a 3’ overhang generated by DNA end resection, mediated by the nucleolytic removal of the complementary strand. The MRE11-RAD50-NBS1 (MRN) complex binds to broken DSBs, and end resection is initiated by a nick generated by the MRE11 nuclease, followed by short range resection being promoted by the CtBP interacting protein (CtIP). Further 5’-3’ resection is mediated by EXO1 and by the combined action of Bloom (BLM) or Werner helicases (WRN) and the DNA2 nuclease (Cannavo and Cejka, 2014; Cejka et al., 2010; Mimitou and Symington, 2008; Niu et al., 2010; Paull and Gellert, 1998; Sartori et al., 2007; Zhu et al., 2008). Extended 3’ ssDNA overhang is coated by ssDNA binding protein RPA. Replacement of RPA by the RAD51 recombinase with the help of mediator proteins (BRCA2, RAD51 paralogs) leads to the formation of a nucleoprotein filament, which is used for homology search and strand invasion to facilitate error free DNA repair by HR (San Filippo et al., 2008; Sugiyama and Kowalczykowski, 2002). In case of extensive end resection across tandem DNA repeats, DSBs may also be mended by the pairing of homologous repeats leading to the loss of interspersed sequences, a process referred to as single strand annealing (SSA) (Bhargava et al., 2016). NHEJ directly ligates broken unresected DSBs, but small deletions may arise when broken DNA ends are degraded. TMEJ acts on partially resected DNAs and relies on the extension of short complementary DNA stretches (microhomology) by POLQ to seal broken DNA, at the expense of losing genetic information (Saito et al., 2017; Wood and Doublie, 2016). The detailed mechanisms determining whether the initially formed 3’ overhangs are further resected for HR or are used as a substrate for error prone TMEJ remain unknown.

The MMR proteins MSH2-MSH3 have also been implicated in HR. MSH2-MSH3 is required for proper ATR-dependent DNA damage signaling to assist HR (Burdova et al., 2015). MSH2-MSH3 helps to reject strand invasion when RAD51-coated single-stranded DNA hybridizes with a not perfectly identical (homoeologous) template DNA (Chen and Jinks-Robertson, 1998; Goldfarb and Alani, 2005; Hum and Jinks-Robertson, 2019; Myung et al., 2001). In addition, MSH2-MSH3 removes secondary DNA structures formed on resected ssDNA. Nevertheless, it remains unclear how the MSH2-MSH3 heterodimer is recruited to DSBs and how it contributes to the downstream steps of HR. A possible connection may be the ‘<u>S</u>WI/SNF-related <u>M</u>atrix-Associated <u>A</u>ctin-Dependent <u>R</u>egulator of <u>C</u>hromatin Subfamily <u>A</u> containing <u>D</u>EAD/H Box1’ protein. It is known to promote the resection of DSB ends in both yeast (Fun30) and human cells (Chen et al., 2012; Costelloe et al., 2012). SMARCAD1 is also known to interact with MSH2-MSH6 for proper MMR (Takeishi et al., 2020; Terui et al., 2018). However, it is still not clear how the interaction between SMARCAD1 and MMR proteins modulates MMR and HR activities. Another protein involved in MMR and HR is EXO1. Although it is known how EXO1 is activated by interaction with MSH2 in MMR, it is not known whether and interaction between EXO1 and MMR may have a role in HR (Schmutte et al., 1998; Tishkoff et al., 1998; Wilson et al., 1998)(Goellner et al., 2018; Schmutte et al., 2001).

The 5’ to 3’ EXO1 exonuclease functions both in MMR and HR (Schmutte et al., 1998; Tishkoff et al., 1998; Wilson et al., 1998). MSH2 interacts with the C-terminal domain of EXO1 via its EXO1 binding domain (Goellner et al., 2018; Schmutte et al., 2001). Although the EXO1 interaction with MSH2 in MMR has been intensively studied, the precise mechanisms of how this interaction regulates DNA end resection during HR need to be uncovered.

Here, initially aiming to uncover a role of MSH2-MSH3 in HR, we reveal that MSH2-MSH3 contributes to HR through the interaction with SMARCAD1 and EXO1. We show that SMARCAD1, MSH2-MSH3, and EXO1 are sequentially recruited to broken DNA to initiate DNA end resection. MSH2-MSH3 prevents POLQ recruitment to broken DNA. Furthermore, MSH2-MSH3 inhibits the sealing of broken DNA ends by TMEJ by blocking POLQ polymerase activity on annealed microhomology sequences carrying DNA mismatch. The blockage of TMEJ facilitates error free HR via more extensive EXO1 dependent end resection.

## Results

### Depletion of MSH2 and MSH3 decreases HR

We previously found that the natural compound, baicalein inhibits MMR (Zhang et al., 2016). Baicalein, is a natural compound derived from *Scutellaria baicalensis*, a herb widely used in Chinese traditional medicine. (Chen et al., 2000; Li-Weber, 2009; Taniguchi et al., 2008). We have recently discovered that, baicalein, triggers the selective killing of DNA mismatch repair (MMR) deficient cancer cells (Zhang et al., 2016). Given the reported interconnection between MMR and DSB repair pathways, we aimed to test if baicalein affects DSB repair. We treated U2OS cells with increasing concentrations of baicalein and measured the frequencies of HR, SSA, NHEJ and MMEJ. Using established reporter essays based on the restoration of GFP expression (Bennardo et al., 2008; Bennardo et al., 2009; Pierce et al., 1999), we observed that the frequencies of HR and SSA were decreased by baicalein treatment in a dose dependent manner. By contrast, NHEJ and MMEJ were not significantly affected (Figure 1A). Since baicalein binds to the MutS complex (Zhang et al., 2016), we hypothesized that MutS may mediate the effect of baicalein on HR and SSA. To test this hypothesis, we depleted the components of the MutSα (MSH2, MSH6) and MutSβ (MSH2, MSH3) complexes and assessed the frequency of HR, SSA, NHEJ, and MMEJ. The depletion of the respective proteins was confirmed by Western blotting (Figure S1A). We observed that knockdown of MSH2 or MSH3 strongly reduced the frequencies of HR and SSA, while knockdown of the MutSα specific subunit MSH6 did not show any effect on HR and SSA (Figure 1B). Consistent with baicalein treatment, depletion of MutS components only showed a small effect on NHEJ and TMEJ (Figure 1B). In summary, the reduction of HR and SSA frequencies conferred by MSH2 or MSH3 depletion in line with previous report (Burdova et al., 2015) correlates with the effect observed of baicalein treatment in this study. Thus, MutSβ (MSH2 and MSH3) but not MutSβ (MSH6), contributes to HR and SSA.

**Figure 1.**
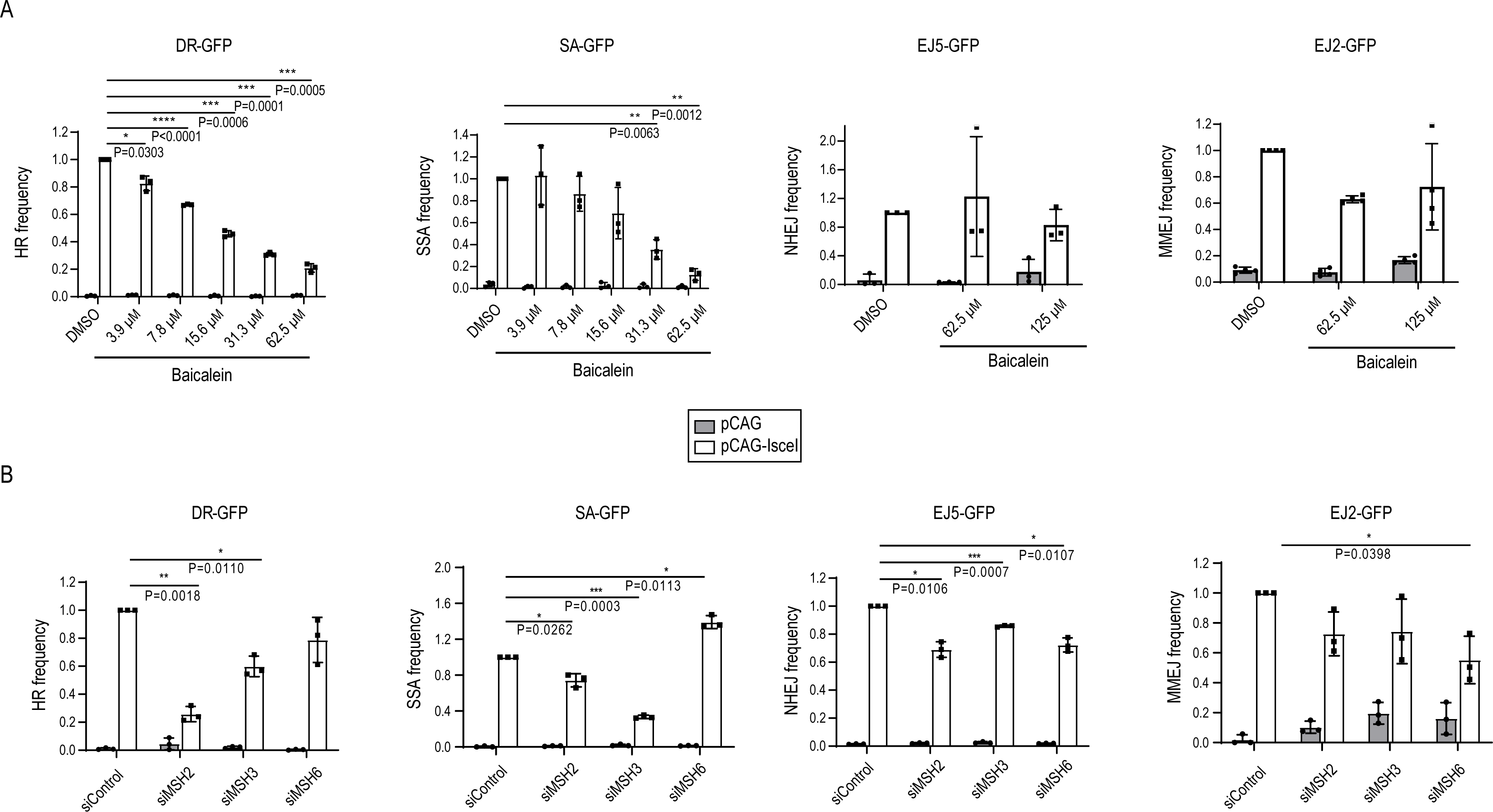
MSH2 and MSH3 functions in homologous recombination (HR) and single strand annealing (SSA). (A). U2OS cells stably expressing DR-GFP, SA-GFP, EJ2-GFP and EJ5-GFP constructs were treated with DMSO or the indicated doses of baicalein for 24 hours and the efficiency of HR, SSA, TMEJ and NHEJ, respectively was determined by scoring and the percentage of GFP positive cells. (B). The efficiency of HR, SSA, TMEJ and NHEJ were measured after transfection of control, MSH2, MSH3 and MSH6 siRNA. Data are represented as mean ± standard deviation (n=3, independent cell culture). P-values were calculated by two-way ANOVA.

### Depletion of MSH2 and MSH3 suppresses DNA end resection

Given that HR and SSA were decreased upon MSH2 or MSH3 depletion, we wanted to determine which step in the HR pathway depends on MSH2 or MSH3. We measured the extent of DNA end resection by assessing RPA2 loading onto resected ssDNA upon treatment with the DSB-inducing agent camptothecin (CPT) by FACS analysis (Forment et al., 2012). We found that RPA2 chromatin binding was reduced upon baicalein treatment (Figure 2A) or MSH2 and MSH3 depletion (Figure 2B). To directly measure the efficiency of the DNA end resection, we used ER-*Asi*SI U2OS cells which allow for generating DSBs via the induction of the *Asi*SI restriction-nuclease upon 4-OHT treatment (Zhou and Paull, 2015). The extent of resection was measured by qPCR assessing amplification from resected ssDNA compared to corresponding dsDNA. Induction of *Asi*SI expression resulted in extensive end resection as measured by the amplification of 335 and 1618 bp fragments, and this end resection was blocked by baicalein treatment (Figure 2C). Consistent with baicalein inhibiting end resection by inhibiting MutSβ (MSH2 and MSH3), end resection was significantly reduced in MSH2 and MSH3 depleted cells. Consistent with our result of HR analysis, no effect was observed upon MSH6 (MutSα) depletion (Figure 2D).

**Figure 2.**
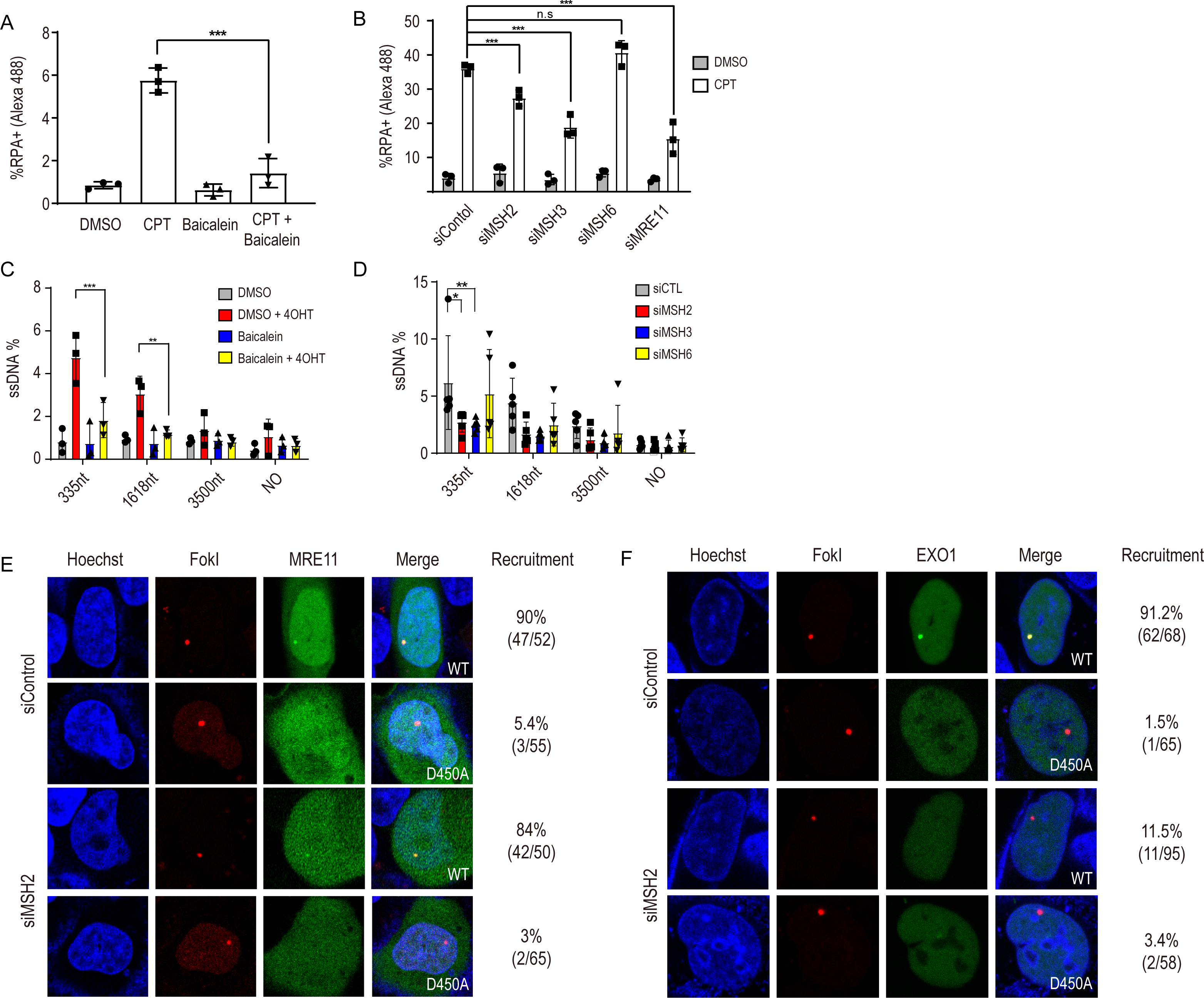
MSH2 and MSH3 are required for DNA end resection. (A) RPA chromatin association was measured after treatment with camptothecin (CPT), baicalein, and the combination of CPT with baicalein. DMSO treatment served as a control. Treatment with baicalein was done for 24 hours at a concentration of 62.5 μM. 5 μ incubation was for 1 hour. Harvested cells were fixed and incubated with RPA2 antibody and the percentage of RPA2 positive cells was analyzed by FACS analysis. (B) RNAi depletion was conducted for 48 hours, and the proportion of RPA2 positive cells was determined by FACS analysis. (C) DSBs were induced in ER-*Asi*SI cells by 4 hours incubation with 4-OHT after treating cells for 24 hours with 62.5 μM of baicalein. Resected DNA was quantified by qPCR after restriction digests (or mock digests) with enzymes cutting double-stranded (but not single-stranded) DNA, 335, 1618 and 3500 nucleotides from the *Asi*SI-induced DSBs. The percentage of amplified ssDNA in relation to DNA amplified from mock treated double-stranded DNA is shown. (D) ER-*Asi*SI cells were transfected with indicated siRNAs and incubated 48 hours. After 4 hours 4-OHT treatment, cells were harvested and subjected to the end resection analyses. (E) mNeon-MRE11 recruitment to *Fok*I induced DSB sites was measured in control and MSH2 knocked down cells. (F) GFP-EXO1 recruitment to *Fok*I induced DSB sites was measured in control cells and upon MSH2 depletion. U2OS cells co-transfected with GFP-EXO1 and *Fok*I WT or the D450A mutant were incubated with Hoechst for 10 min to visualize the nucleus. Cells were then incubated in CO_2_ independent media and live cell images were taken with a confocal microscope. EXO1 recruitment to DSB sites was quantified (right column) by calculating the proportion of cells showing a colocalization of GFP-EXO1 fusion at the DSB site demarcated by the mCherry fusion.

### MSH2-MSH3 interacts with EXO1 to promote end resection activity

We next wished to determine the step of DNA end resection facilitated by MSH2-MSH3 and therefore monitored the recruitment of MRE11 and EXO1 to DSB sites. DSB can be induced and visualized using a fusion protein comprised of *Fok*I, the lac repressor and mCherry, where breaks can be targeted to lac operator repeats in U2OS cells (Shanbhag et al., 2010). Using this assay, we found that MRE11 recruitment to the DSB was not affected in control siRNA and MSH2 siRNA treated cells (Figure 2E) suggesting that MSH2 is not essential for the recruitment of MRE11 to DSBs. By contrast, EXO1 recruitment to DSB sites was significantly decreased in MSH2 (Figure 2F) or MSH3 knockdown cells (Figure S1B). MRE11 and EXO1 recruitment depended on DSB formation as recruitment was not observed when a catalytically inactive *Fok*I D450A nuclease was used (Figures 2E-F and S1B). Collectively, the MSH2-MSH3 complex is required for recruiting EXO1 but not MRE11 to DSBs.

It has been shown that EXO1 interacts with MSH2 through the C-terminal region of EXO1 (Schmutte et al., 2001). We first confirmed this interaction by immunoprecipitation (IP) of the endogenous proteins. As previously observed, MSH2 and EXO1 interact with each other (Figure 3A). This interaction is not changed by ionizing radiation (IR) (Figure 3A). Using a series of GFP tagged EXO1 deletion mutants (EXO1 D1 to D4) spanning the entire protein and myc tagged MSH2, we confirmed that the C-terminal residues 600-846 of EXO1 are needed for MSH2 binding (Figure S2A) (Schmutte et al., 2001). To further investigate how MSH2 regulates the recruitment of EXO1 to DSB sites, we used a series of small C-terminal deletions and narrow down the minimal MSH2 binding domain of EXO1 to aa 776 to 807 (Figure 3B). Conversely, the minimal MSH2 domain required for EXO1 binding domain was determined to be aa 306-623 (Figure S2B, Figure 3C). This domain contains the MutS core domain and is part of previously annotated EXO1 binding domain (Schmutte et al., 2001). The requirement of the EXO1 C-terminal residues 706-807 for MSH2 binding *in vivo* was further confirmed by Cell-based Unidentified Protein Interaction Discovery (CUPID) assays (Lee et al., 2011). For this assay, a PKC-δ domain was fused to mRFP-MSH2 and employed to tether the fusion protein to the nuclear membrane upon PMA (phorbol 12-myristate 13-acetate) treatment, which resulted in the localization of EXO1, but not the EXO1 D13 (Δ792-807) to the nuclear membrane (Figure S2C).

**Figure 3.**
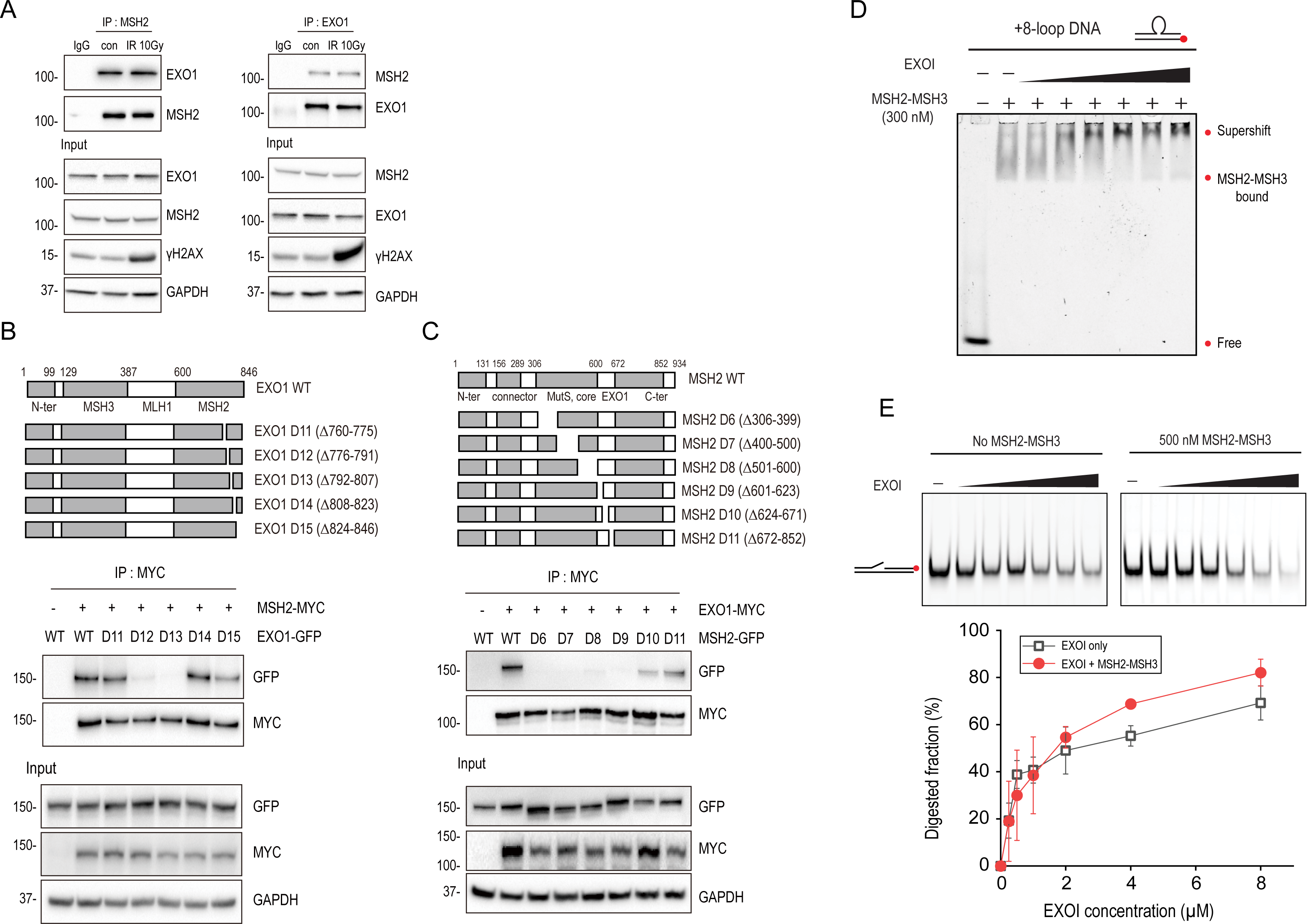
MSH2 interaction with EXO1 enhances the EXO1 activity. (A) For endogenous immunoprecipitation, 0 or 10 Gy of ionizing radiation were irradiated to HEK293T cells. Extracts were incubated with anti-IgG, anti-MSH2 or anti-EXO1 antibody and immunoprecipitated proteins by dynabeads protein G were analyzed by western blot analysis. γH2AX was used as a DNA damage marker for 10 Gy IR exposure. (B) Diagram of EXO1 wild type and each deletion mutant. Each GFP-EXO1 and myc-MSH2 WT was co-transfected to HEK293T cells and interaction of each EXO1 deletion mutant with MSH2 was determined by the MYC antibody immunoprecipitation. (C) Diagram of MSH2 WT and each deletion mutant. HEK293T cells were co-transfected with myc-EXO1 WT and each GFP-MSH2 deletion mutant and interaction of each MSH2 deletion mutant with EXO1 was determined by the MYC antibody immunoprecipitation. (D) EMSA for MSH2-MSH3 and EXO1. 300 nM MSH2-MSH3 was bound to +8-loop DNA, and EXO1 was then titrated (0, 50, 100, 200, 400, 800, and 1600 nM). (E) Nuclease activity of EXO1 in the presence or absence of MSH2-MSH3. 20 nM DNA with a 90 bp flap DNA was reacted with EXO1 at different concentrations (0, 250, 500, 1000, 2000, 4000, 8000 nM) in the absence (top left) or presence (top right) of 500 nM MSH2-MSH3. Experiments were performed in triplicate and quantified (bottom). Error bars were obtained from standard deviation.

To investigate the direct interactions between MSH2-MSH3 and EXO1 in vitro, all proteins were purified (Figure S2D), and their activities of the purified proteins were tested (Figure S2E). Consistent with the previous work, MSH2-MSH3 showed higher binding affinity to an oligonucleotide substrate carrying an 8-nt loop (+8-loop DNA) compared to a corresponding homoduplex oligonucleotide substrate (Figure S2E) (Kumar et al., 2014). In addition, EXO1 displayed the expected nuclease activity when served with a 40 bp DNA double-stranded substrate carrying a 4-nt single-stranded 3’ overhang (Figure S2F). However, EXO1 did not show high exonuclease activity for blunt end DNA compared to 3’ overhang DNA (Figure S2F).

Having confirmed the functionality of *in vitro* purified MSH2-MSH3 and EXO1 proteins, we asked if MSH2-MSH3 is able to facilitate recruiting of EXO1 to DNA substrates. We performed supershift assays adding increasing concentrations of EXO1 to MSH2-MSH3 bound to an +8-loop-containing oligonucleotide substrate and observed a dose dependent supershift (Figure 3D). The supershift by EXO1 was observed at lower concentration (∼100 nM) in the presence of MSH2-MSH3, while EXO1 alone bound to the same DNA substrate only at a much higher concentration (∼800 nM), indicating that MSH2-MSH3 promotes the association of EXO1 with DNA (Figure 3D, S2G).

We then asked how the interaction between MSH2-MSH3 and EXO1 contributes to DNA end resection. It was reported that MSH2-MSH6 enhances EXO1 activity in MMR (Genschel and Modrich, 2003). Thus, we tested whether MSH2-MSH3 also enhances the nuclease activity of EXO1. In the presence of MSH2-MSH3, DNA degradation activity of EXO1 increased, suggesting that the recruitment of EXO1 by MSH2-MSH3 enhances the end resection (Figure 3E).

### SMARCAD1 directly interacts to MSH2-MSH3

MSH2-MSH3 preferentially recognizes small loop structures. Since resected DSBs do not have small loops, we asked how MSH2-MSH3 might be recruited to DSB sites to facilitate EXO1 recruitment for end-resection. We hypothesized that protein(s) interacting with MSH2-MSH3 help recruit MSH2-MSH3 to DSBs. The chromatin remodeler SMARCAD1 is known to interact with MSH2-MSH6 (Takeishi et al., 2020; Terui et al., 2018) and has been reported to be recruited to DSBs to facilitate the end resection in yeast and human cells (Chakraborty et al., 2018; Chen et al., 2012). To test if SMARCAD1 also recruits MSH2-MSH3, we first re-examined if SMARCAD1 interacts with MSH2 in HEK293T cells. We observed a reciprocal pull down of endogenous MSH2 and SMARCAD1 using IP of the endogenous proteins (Figure 4A-C). Similar to the interaction between MSH2 and EXO1, the interaction between SMARCAD1 and MSH2 was not changed by IR treatment. To identify the MSH2 binding domain in SMARCAD1, we generated GFP tagged SMARCAD1 WT and a series of deletion mutants (D1 to D4) spanning the entire protein and assessed association with myc-MSH2 by IP with myc antibodies. Based on bioinformatic analysis, it was predicted that SMARCAD1 has a potential MSH2 binding domain, termed SHIP box, in its N-terminus (aa residues 5-11) (Goellner et al., 2018). Our domain analysis experimentally confirmed that the N-terminus of SMARCAD1 is required for MSH2 binding, with SMARCAD1 D1 (Δ1-156) deletion being unable to bind MSH2 (Figure 4B).. Conversely, in co-transfection experiments with a series of deletion mutations in MSH2,, we narrowed the minimal SMARCAD1 binding domain in MSH2 to aa residues 306 to 623 (Figure S3A, Figure 4C) which were the same region of MSH2 interacting with EXO1 (Figure 3C), suggesting SMARCAD1 and EXO1 bind to the same region of MSH2. Although MSH2 D6 to D9 deletion mutants spanning aa 306 to 623 lost interaction with EXO1 and SMARCAD1, these MSH2 deletion mutants were still able to assemble a MutSβ heterodimer complex with MSH3 (Figure S3B). We tested the interaction between SMARCAD1 and EXO1 by endogenous IP, but we were unable to detect an interaction under these conditions (Figure S3C). The interaction between MSH2 and SMARCAD1 *in vivo* was confirmed by CUPID analysis (as described above) (Figure S3D).

**Figure 4.**
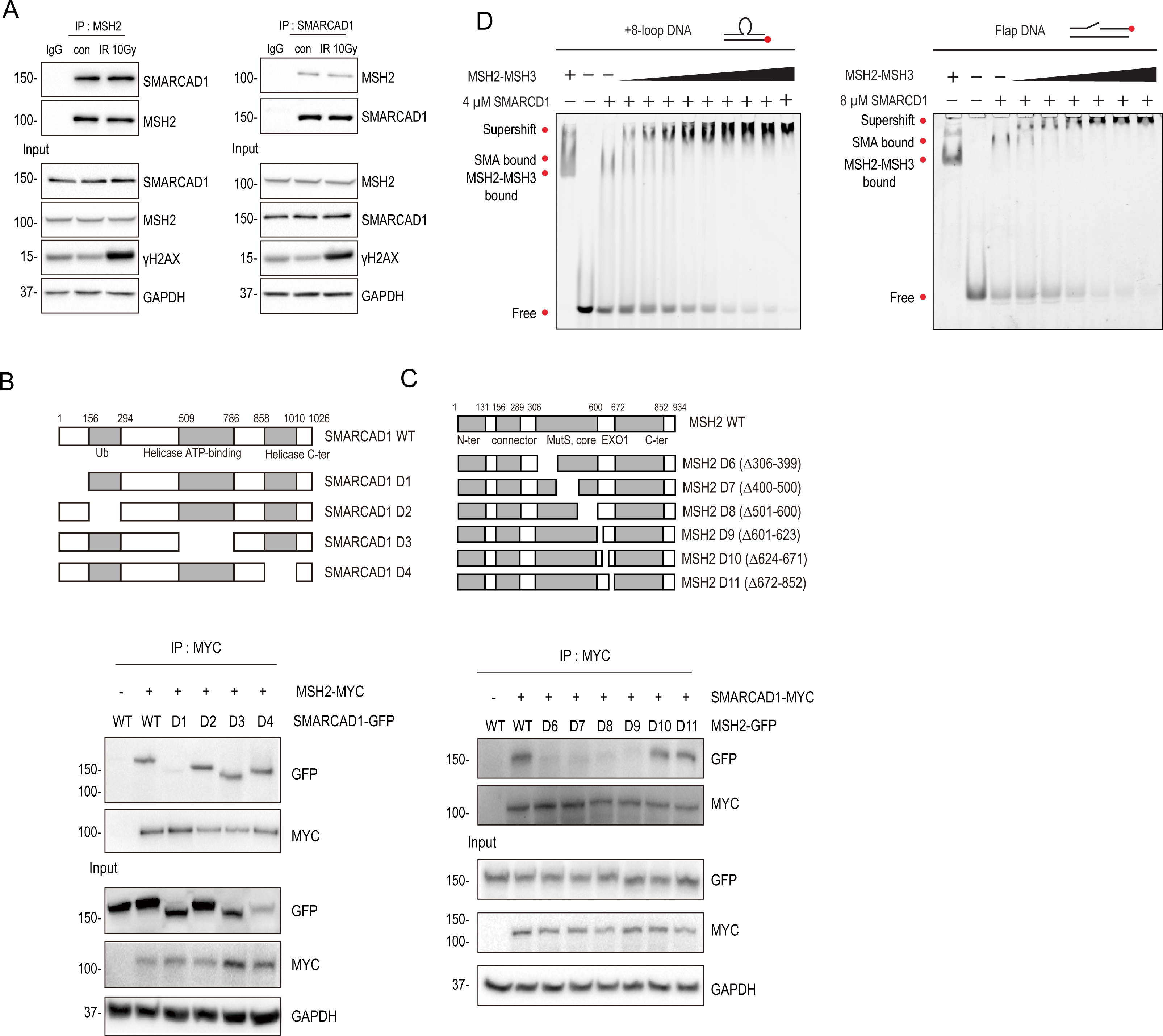
SMARCAD1 directly interacts with MSH2. (A) 0 or 10 Gy of IR were irradiated to HEK293T cells. Extracts were immunoprecipitated with IgG, MSH2 or SMARCAD1 antibody and immunoprecipitated proteins by dynabeads protein G were analyzed by western blot analysis. γH2AX was used as a DNA damage marker for 10 Gy IR exposure. (B) Diagram showed SMARCAD1 WT and each deletion mutant. Cells were co-transfected with myc-MSH2 WT and GFP-SMARCAD1 WT and each mutant and interaction of each deletion mutant of SMARCAD1 with MSH2 was determined by the MYC antibody immunoprecipitation. (C) MSH2 WT and each deletion mutants were shown. Myc-SMARCAD1 WT was co-transfected with each GFP-MSH2 WT and each mutant and interaction of each deletion mutant of MSH2 with SMARCAD1 was determined by the MYC antibody immunoprecipitation. (D) Electrophoretic mobility shift assay (EMSA) μM SMARCAD1 was bound to +8-loop DNA, and then MSH2-MSH3 was titrated (0, 10, 20, 40, 80, 100, 150, 200, 300, and 400 nM, left). 8 μM SMARCAD1 was bound to 58 bp flap DNA, and then MSH2-MSH3 was titrated (0, 10, 20, 40, 80, 100 and 150 nM, right).

To determine if SMARCAD1 directly interacts with MSH2-MSH3, we performed *in vitro* binding assays with purified proteins. After SMARCAD1 was incubated with the +8-loop DNA, MSH2-MSH3 was added at different concentrations. As MSH2-MSH3 concentration increased, the SMARCAD1-DNA band progressively disappeared and a supershifted band emerged (Figure 4D). The supershifted band by MSH2-MSH3 started to appear from 10 nM concentration, whereas the binding of MSH2-MSH3 alone to +8-loop DNA occurred from 80 nM (Figure S2E). We also tested another DNA construct, flap DNA. The flap DNA structure is much similar to the sequences exist in the DSB. SMARCAD1 was pre-incubated with the flap DNA and then MSH2-MSH3 was titrated. MSH2-MSH3 bound to the flap DNA at lower concentration with SMARCAD1 (10 nM of MSH2-MSH3) than without SMARCAD1 (80 nM of MSH2-MSH3) (Figure 4D and Figure S3E). Our results demonstrate the binding affinity of MSH2-MSH3 to +8-loop and flap DNA is ∼ 8 times enhanced in the presence of SMARCAD1 and support that SMARCAD1 recruits MSH2-MSH3 and forms a complex on DNA.

### Interdependence of SMARCAD1, MSH2 and EXO1 location at DNA damage sites

To determine the interdependence of SMARCAD1, MSH2 and EXO1 recruitment to DNA damage sites, GFP-tagged versions of these genes were transfected and their recruitment to microirradiated strips was determined. We found that GFP-MSH2 accumulated at microirradiation sites; accumulation being compromised upon SMARCAD1-, but not EXO1 depletion (Figure 5A). Conversely, SMARCAD1 recruitment was not altered by MSH2 and EXO1 depletion (Figure 5B). Finally, EXO1 recruitment depended on both MSH2 and SMARCAD1 (Figure 5C). Taken together, these data indicate that SMARCAD1 recruits MSH2 and then MSH2 recruits EXO1 to DSB sites.

**Figure 5.**
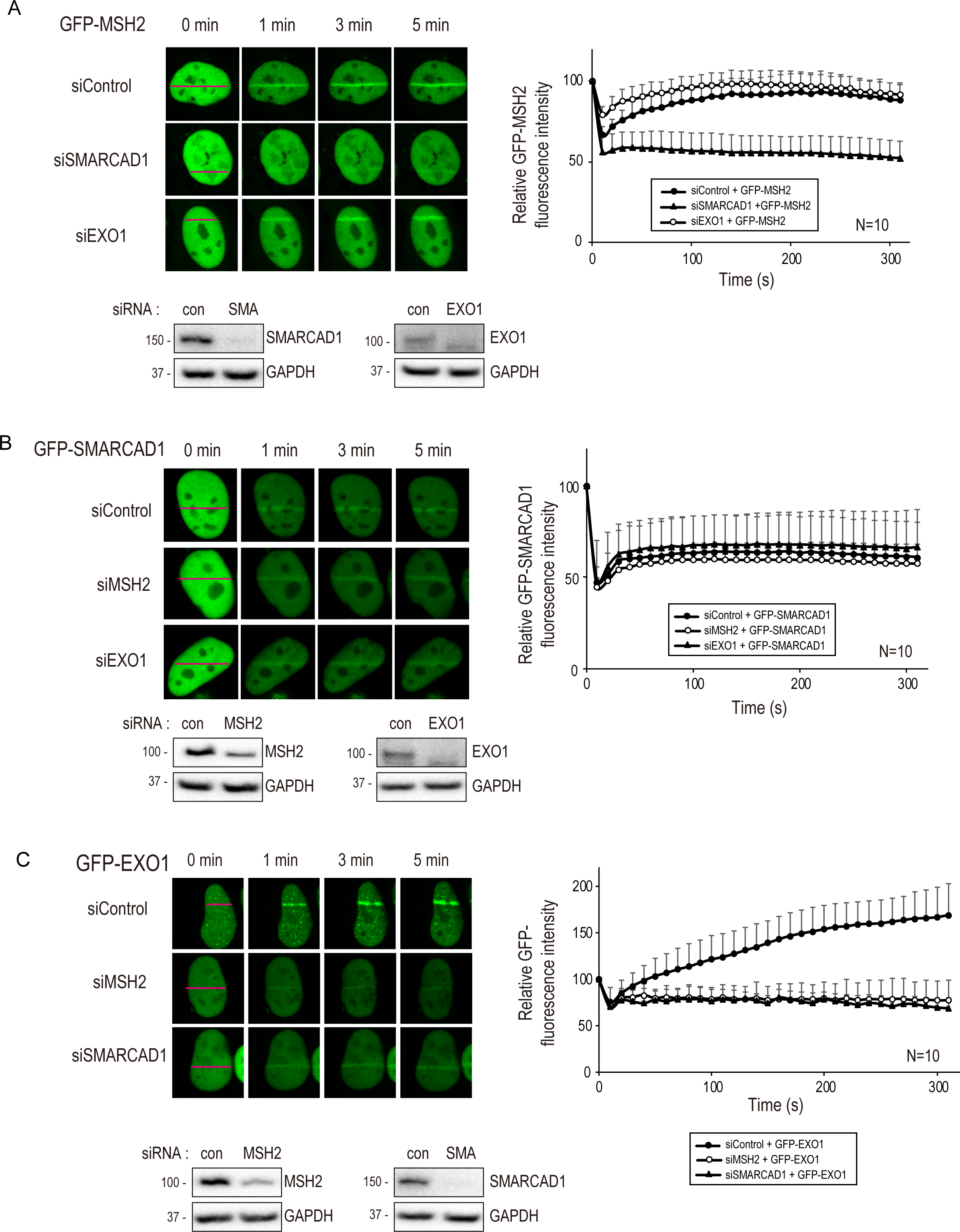
SMARCAD1, MSH2 and EXO1 move to DSB sites in orderly manner.(A, B, C) U2OS cells were transfected with control or indicated siRNA. GFP-MSH2 (A), GFP-SMARCAD1 (B), and GFP-EXO1 (C) were transfected after 24 hours and cells were 10 μM of BrdU for 24 hours. Cell images were taken after microirradiation in 10 seconds interval for 5min with confocal microscopy.

Since MRE11 recruitment to DSB sites was not dependent on MSH2 (Figure 2E), we examined the interaction between MRE11 and SMARCAD1. We found that MRE11 recruitment to the microirradiation induced DSB sites occurred normally in both control, MSH2 and SMARCAD1 knockdown cells (Figure S4A) indicating that MRE11 recruitment to DSB sites is independent from SMARCAD1 and MSH2 recruitment. MMR proteins have been suggested to function to reject heteroduplex DNA with imperfect matches during the later stages of the HR (Chen and Jinks-Robertson, 1998; Goldfarb and Alani, 2005; Hum and Jinks-Robertson, 2019; Myung et al., 2001). The formation of heteroduplex DNA requires RAD51 dependent strand invasion. Thus, we asked if MSH2-MSH3 protein recruitment to DSB sites depends on RAD51. To validate whether RAD51 affects MSH2 recruitment to DSB sites, we treated cells with the RAD51 inhibitor B02 for 4 hours and monitored the MSH2 recruitment to the microirradiation induced DSB sites. MSH2 accumulation to the microirradiation induced DSB was not affected by the RAD51 inhibitor treatment (Figure S4B), indicating that MSH2 is upstream of RAD51 and recruited before the strand invasion stage of the HR. To investigate if MMR proteins unrelated to MSH2-MSH3 are involved in EXO1 recruitment to DSB sites, we knocked down MLH1, the downstream factor of MutS, but did not find altered EXO1 recruitment (Figure S4C).

### SMARCAD1-MSH2-EXO1 recruitment is important for HR

We next assessed the requirement of various SMARCAD1 domains for recruitment to microirradiated sites. The SMARCAD1 D1 deletion mutant, which lost interaction with MSH2, was well recruited to the DNA damage sites like wild type SMARCAD1 (Figure 6A). Interestingly the recruitment of SMARCAD1 D3 deletion mutant, which lacks the DNA helicase and ATP-binding domain, was decreased, suggesting that DNA helicase activity is important for DNA binding (Figure 6A). To directly test if SMARCAD1 is required for HR we conducted reporter-based HR assays (see above) and found that SMARCAD1 depletion decreased HR efficiency (Figure 6B). The reduction of HR is due to the specific SMARCAD1 depletion. HR being restored upon transfection with the RNAi resistant full length SMARCAD1, but not by the siRNA resistant SMARCAD1 D1 mutant (Figure 6B). Consistent with the results of the DR-GFP assay, the SMARCAD1 D1 mutant could not rescue RAD51 foci formation after IR 10 Gy treatment unlike the SMARCAD1 WT in siRNA knockdown cells (Figure 6C). These data show that SMARCAD1-dependent MSH2 recruitment is important to promote HR.

**Figure 6.**
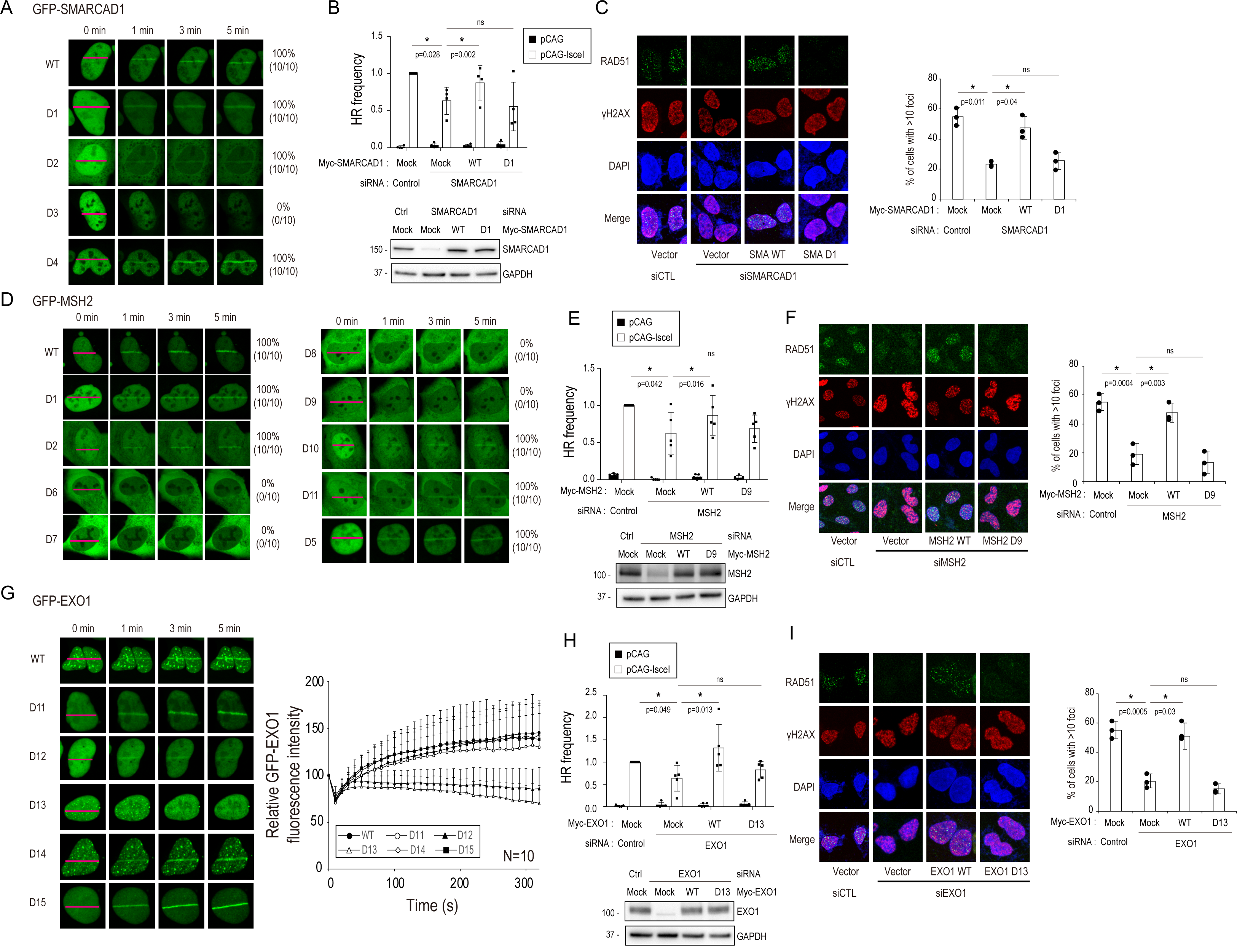
SMARCAD1, MSH2 and EXO1 are required for HR. (A) U2OS cells transfected with GFP-SMARCAD1 WT or each deletion mutant were microirradiated (MI) and their movements to MI-induced DSB was monitored under confocal microscopy after incubation with 10 μM of BrdU for 24 hours. (B) HR frequency was determined in SMARCAD1 knocked down DR-GFP U2OS cells after transfecting either myc-SMARCAD1 WT or D1 deletion mutant. (C) SMARCAD1 knocked down U2OS cells after transfecting either myc-SMARCAD1 WT or D1 deletion mutant were irradiated with 10 Gy IR and RAD51 foci in the nucleus were counted under confocal microscopy. (D) U2OS cells transfected with GFP-MSH2 WT or each deletion mutant was monitored under confocal microscopy after incubation with 10 μM of BrdU for 24 hours. (E) HR frequency was determined in MSH2 knocked down DR-GFP U2OS cells after transfecting either myc-MSH2 WT or D9 deletion mutant. (F) MSH2 knocked down U2OS cells after transfecting either myc-MSH2 WT or D9 deletion mutant were irradiated with 10 Gy IR and RAD51 foci 51 in the nucleus were counted under confocal microscopy. (G) U2OS cells transfected with GFP-EXO1 WT or each deletion mutant was monitored under confocal microscopy after incubation with 10 μM of BrdU for 24 hours. (H) HR frequency was determined in EXO1 knocked down DR-GFP U2OS cells after transfecting either myc-EXO1 WT or D13 deletion mutant. (I) EXO1 knocked down U2OS cells after transfecting either myc-EXO1 WT or D13 deletion mutant were irradiated with 10 Gy IR and RAD51 foci in the nucleus were counted under confocal microscopy.

We next examined the recruitment of wild type GFP-MSH2 and a series of GFP-MSH2 deletion mutants that include mutants D6 to D9 spanning the aa 306 to 623 of MSH2 required for SMARCAD1 and EXO1 binding (Figure 4C). We found that wild type MSH2, but not the EXO1 and SMARCAD1 binding defective D6 to D9 mutants were recruited to microirradiated strips (Figure 6D). To exclude the possibility that D6 to D9 MSH2 fail to be recruited to microirradiated strips because of not entering the nucleus, MSH2 D6 to D9 deletion constructs were tagged with a nuclear localization signal (NLS). Forcing MSH2 D6 to D9 to the nucleus failed to affect recruitment to microirradiated stripes. (Figure S5A). Thus, MSH2 recruitment to DNA damage sites depends on SMARCAD1. Measuring the effect of MSH2 depletion on HR activity, and RAD51 foci formation upon IR treatment, we confirmed that HR is reduced by MSH2 depletion (Figure 6E) and found that the MSH2 depletion can be rescued by expressing siRNA resistant full length MSH2, but not by the SMARCAD1 binding defective MSH2 D9 mutant (Figure 6E, F). These results suggest that MSH2 recruitment to DNA damage sites depends on SMARCAD1 and this recruitment is important for proper HR.

Lastly, to determine if EXO1 recruitment to DSB sites is affected by MSH2 binding activity, we asked if GFP-EXO1 and GFP-EXO1 D11 to 15 deletion mutants, the D12 (aa776-791) and D13 (aa 792-807) mutants being defective for MSH2 binding (Figure 3B), are recruited to microirradiated strips and facilitate HR and RAD51 foci formation (Figure 6G, H, I). We found that wild type GFP-EXO1 and all GFP-EXO1 mutants except for D12 and D13 were recruited to microirradiated sites (Figure 6G), and that the MSH2 interaction domain of EXO1 (tested by the D13 deletion) is required for efficient HR (Figure 6H) and RAD51 foci formation (Figure 6I). Conversely, the decreased EXO1 recruitment to microirradiated stripes in MSH2 depleted cells were rescued by the transfection with siRNA resistant wild type MSH2, but not by the MSH2 D9 (as 601-623) deletion mutant which is defective for EXO1 binding (Figure S5B). These data show that EXO1 recruitment to damage sites requires MSH2 binding. Collectively our data show that EXO1 is required to damage sites by the MSH2-MSH3 complex, which in turn is targeted by SMARCAD1.

To exclude the possibility that any of the RNAi depletion experiments alter the cell cycle profile, which in turn would affect HR activity, we monitored cell cycle profiles by FACS analysis and confirmed that the depletion of MSH2, MSH3, SMARCAD1 or EXO1 did not change the cell cycle profile (Figure S5C).

### MSH2 inhibits POLQ-mediated end-joining

It is known that heteroduplex DNAs are rejected by mismatch repair proteins during HR and SSA (Sugawara et al., 2004). More recently, DNA polymerase θ (POLQ)-mediated end-joining (TMEJ) was shown to use the pairing of short homologous sequences (2–6 bp, microhomology) of resected ssDNA to prime and extend DNA synthesis to mend double strand breaks, at the cost of generating small deletions. We postulated that MSH2-MSH3 might act on resected DNA to prevent POLQ recruitment, allowing for further DNA end resection, facilitating error free HR. Such mechanism would require POLQ being able to prime from mismatch containing heteroduplex DNA. Although we did not see drastic effect on MMEJ after MSH2 knockdown (Figure 1B), which would be due to the sensitivity of assay. It is possible that TMEJ activity is covered by NHEJ, which can also perform MMEJ. We decided to study the relationship between POLQ and MSH2-MSH3 more directly. We therefore tested if POLQ can extend primer carrying a 2 bp mismatch at different positions (Figure S6) and found that this is the case. In a control experiment, the polymerase fragment of POLQ catalyzed template-dependent DNA synthesis from a perfectly annealed primer pair similar to the exonuclease-deficient *E. coli* pol I Klenow fragment (Kf exo-, another A-family DNA polymerase which served as a control (Figure S6A) (Hogg et al., 2011; Yousefzadeh et al., 2014). Importantly, annealed primer pairs carrying 2 bp mismatches, 1 and 2 bp or 3 and 4 bp upstream from the 3’ primer end (MM-1, 2 and MM-3, 4) could be much more efficiently extended by POLQ as compared to Kf exo- (Figures S6B and S6C), an effect particularly strong when the mismatch occurs at the primer junction (Figure S6B) or 2 bases downstream (Figure S6C). Having shown that POLQ can extend DNA synthesis from mismatched primers, we chose the MM-3, 4 substrate (Figure S6C) to test the effect of MSH2-MSH3 on POLQ activity. POLQ activity was inhibited by MSH2-MSH3 only in the presence of a 2 bp mismatch in the double-stranded primer (Figures 7A-C), compared to perfectly matched primers (Figures 7D-F). The termination probabilities at N3 position were significantly increased in the presence of MSH2-MSH3 when the 2bp-mismatched substrates were used (Figures 7A and B). In consistent with these results, the amount of fully extended products was reduced in the reactions with the 2 bp-mismatched substrates (Figure 7C), but not with no-mismatch substrates (Figures 7D-F). MSH2-MSH3 with MSH2 G674A inhibited the POLQ activity more strongly than wild-type MSH2-MSH3 (Figures. S6E-G). MSH2 G674A is defective in ATP binding-induced dissociation from mismatched DNA substrates (Figure S2E) (Geng et al., 2012). Our results indicate that MSH2-MSH3 can inhibit POLQ activity when priming from mismatched DNA substrates. To substantiate the importance of mismatched DNA substrates for DNA extension in vivo, and the role of MSH2-MSH3 in preventing this, we assessed if POLQ recruitment to DNA damage sites is affected by MSH2 depletion, and found that this is the case. POLQ is recruited to damage sites and this is strongly increased upon MSH2 depletion (Figure 7G). Mutation signatures in the genome can predict how DNA damage is repaired by different DNA repair pathway. The error-prone TMEJ-dependent repair often results in small deletion/insertion with microhomology signature at the breakpoint junction (Hwang et al., 2020). When the CEL locus was cleaved by CRISPR-Cas9, MSH2 knockdown increased the frequency of deletion with microhomology at the breakpoint junctions (Figure 7H). Collectively, MSH2 and MSH3 are recruited to DSB sites by SMARCAD1 to promote DNA end resection by recruiting EXO1 to facilitate error free HR. This process inhibits POLQ recruitment and TMEJ at DSB sites.

**Figure 7.**
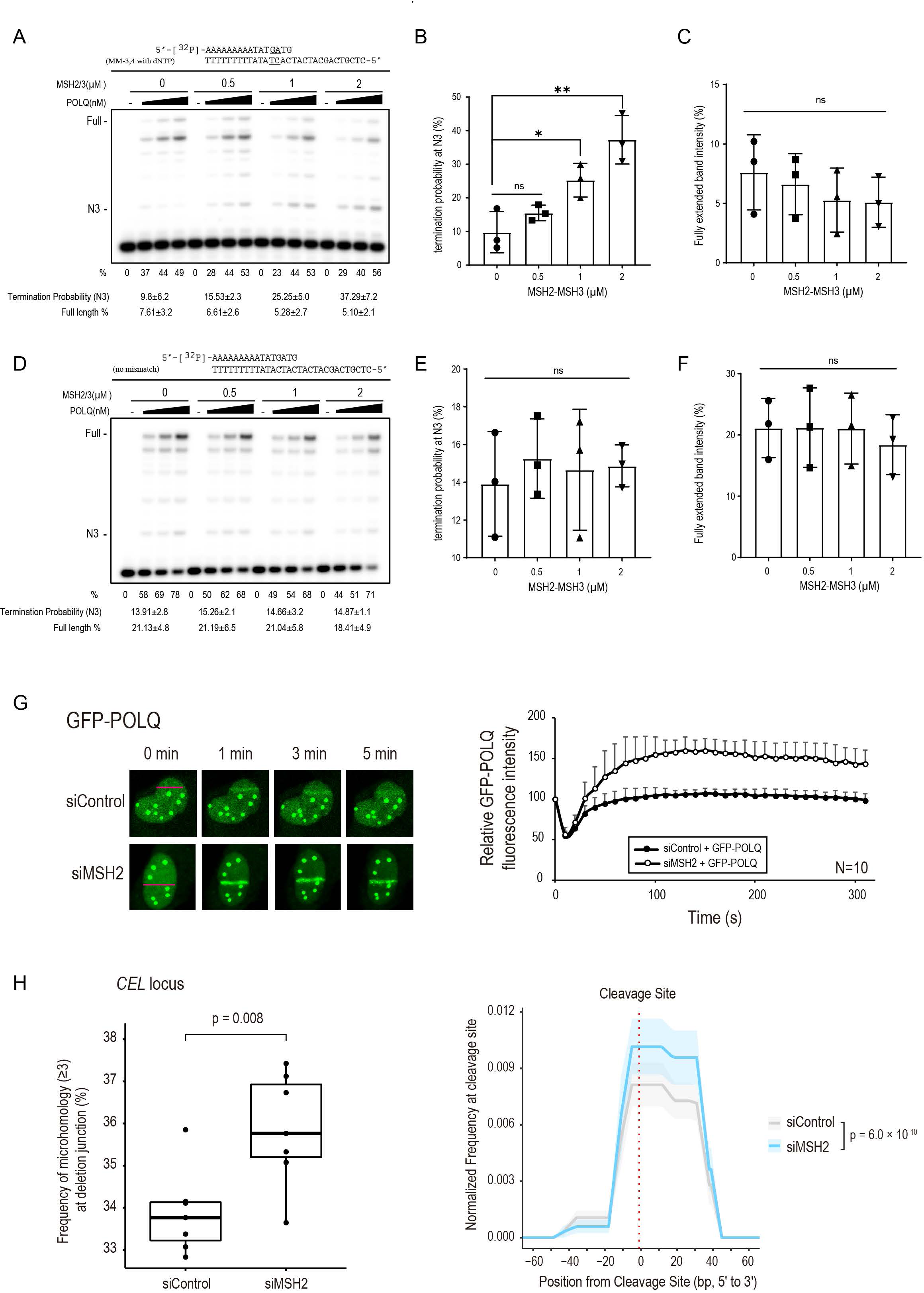
MSH2-MSH3 inhibits POLQ extension from a mismatched primer. Increasing amounts of POLQ (0.3, 0.6, 1.3 nM) were incubated in the presence of indicated amount of MSH2-MSH3 with the 5′-^32^P-labeled primer templates described on top of the gel in the presence of all 4 nt at 37°C for 10 minutes. The first lane (-) contained no enzyme. (A) The percentage (%) of the product extension from the primer is shown below each lane. 2 mismatched base pairs were placed at the 3^rd^ and 4^th^ bp from the primer-template junction. (B) The termination probability at position N3 is defined as the band density at N3 divided by the intensity of ≥ N3. (C) The amount of full-length extension products is defined as the fully extended band density divided by the intensity of ≥ N0 (Primer position). The effects of MSH2-MSH3 on non-mismatched substrate are measured similarly (D, E, F). (G) POLQ movements to MI induced DSB sites were monitored in U2OS cells transfected with control or MSH2 siRNA after incubation with 10 μM of BrdU. (H) Mutation signatures at the CEL locus upon CRISPR-Cas9 induced DSB were compared in control and MSH2 knockdown HEK293T cells. Boxplot showing frequency of deletion mutations harboring microhomology longer than two nucleotides at the DNA junction out of total deletion mutations induced by CRISPR-Cas9 targeting CEL locus in control and MSH2 knockdown HEK293T cells. P-value was calculated by unpaired two-tailed t-test (n=7) (left). DNA deletion spectrum associated microhomology longer than 4 nucleotides induced by CRISPR-Cas9 targeting CEL locus in control and MSH2 knockdown HEK293T cells. P-value was calculated by paired two-tailed t-test (n=7) (right).

## Discussion

In summary we show that the MSH2-MSH3 promotes error free DSB repair by facilitating HR by two complementary mechanisms: MSH2-MSH3 is recruited after the initial stages of DNA end resection by SMARCAD1. SMARCAD1 and MSH2-MSH3 dependent recruitment of the EXO1 nuclease allows for further resection, promoting error free HR. At the same time, the MSH2-MSH3 complex also acts in cis, by inhibiting the access of POLQ, preventing POLQ priming and extension from mismatched DNA, that results from imperfect pairing of partially resected DNA ends.

The involvement of MSH2-MSH3 in the HR has been previously suggested. Based on their activity to recognize mismatched DNA sequences, MSH2-MSH3 has been discussed to function at the later, postsynaptic stage of the HR, helping to reject invading strands that do not perfectly match template DNA (Burdova et al., 2015; Chen and Jinks-Robertson, 1998; Goldfarb and Alani, 2005; Hum and Jinks-Robertson, 2019; Myung et al., 2001). In addition, several studies implemented MSH2-MSH3 proteins function at the early stages of the HR. MSH2-MSH3 were discussed to inhibit hairpin structures forming during DNA end resection. Moreover, MSH2-MSH3 was discussed to facilitate full DNA damage checkpoint activation (Burdova et al., 2015).

However, none of these studies showed a direct role of MSH2-MSH3 for DNA end resection. The sequential recruitment of SMARCAD1, MSH2-MSH3, and EXO1 observed in the present study, together with requirement of these proteins for RAD51 loading and proper HR clearly demonstrates that MSH2-MSH3 have an active role at an early stage of the HR. In addition to inhibiting TMEJ by rejecting POLQ, MSH2-MSH3 recruitment to DSB sites facilitates EXO1 recruitment and long-range DNA end resection, thus funneling pathway choice towards error free HR. Besides facilitating EXO1 recruitment, how could MSH2-MSH3 further aid EXO1 exonucleolytic activity for DNA end resection? When MSH2-MSH3 binds to loop structures, DNA bound by MSH2-MSH3 is bent for proper recognition by downstream proteins (Gupta et al., 2011). It is thus possible that MSH2-MSH3 recruited to DSB sites could bend DNA to provide for a better access of EXO1 to its DNA substrate. When DSBs are repaired, as part of the early stages of DNA end resection, MRE11-RAD50-NBS1 generates a nick and degrades ssDNA with a 3’ to 5’ polarity (Chapman et al., 2012). As part of this reaction, small single-stranded gaps arise, and these structurally resemble small loop structures, the single-stranded DNA stretch being extruded. MSH2-MSH3 would recognize such a structure (Kumar et al., 2014) and bend this small gapped DNA to provide for an entry platform for the EXO1 5’ to 3’ exonucleolytic reaction needed for generating long ssDNA.

Chromatin remodeling complexes play important roles in DSB repair. During DSB repair SMARCAD1 is known to be phosphorylated by ATM and ubiquitinated by RING1, both modifications being important for DNA end resection (Chakraborty et al., 2018). The budding yeast Fun30 protein, the ortholog of SMARCAD1 is a major nucleosome remodeler enhancing Exo1 and Sgs1 dependent end resection during HR repair (Chen et al., 2012). Mammalian SMARCAD1 was equally suggested to have a role in HR in mammals (Chakraborty et al., 2018). Moreover, a role for Fun30 for MMR through its interaction with MSH2 was described (Goellner et al., 2018; Terui et al., 2018). However how SMARCAD1 promotes the end resection for the HR was not well established. Exploring the requirement of SMARCAD1 for MSH2-MSH3 targeting was motivated by speculating that SMARCAD1 is a chromatin remodeler that may unwind chromatin structures near the DSB sites and help recruiting MSH2-MSH3 to DSB sites at the early stages of DSB processing. Given that interactions between SMARCAD1 and MSH2, and between MSH2 and EXO1 are conserved, it appears that these interactions are equally used to facilitate HR throughout evolution. Our results clearly support a conserved mechanism where SMARCAD1 interacts with MSH2-MSH3 and enhances the DNA binding affinity of MSH2-MSH3. (Figure 4D). As part of this process, MSH2-MSH3 prevents the access of POLQ, the key enzyme facilitating error-prone TMEJ. Analogous to its role in the MMR reaction (Goellner et al., 2018), MSH2-MSH3 recruits EXO1 for promoting DNA end resection. Interestingly, the MSH2 domains needed for SMARCAD1 and EXO1 are placed in the same part of MSH2. Albeit we did not observe competition between SMARCAD1 and EXO1 for MSH2 binding in overexpression experiments (data not shown), it is possible that SMARCAD1 may lose its interaction with MSH2 allowing EXO1 to occupy the same region of MSH2 to promote the end resection.

From the evolutionary point of view, it makes sense that key modalities such as the MSH2-MSH3 dependent recruitment of EXO1 are used by more than one repair modality. Shared modalities might also facilitate the crosstalk between pathways, to allow for more efficient repair. Especially in case of HR, where a single persistent DSB might lead to lethality, cross talks might equally help to ensure that all lesions are mended. We consider it likely that MSH2-MSH3 has an important role in preventing error prone repair by POLQ. However, detecting such an effect, imprinted in mutational signatures associated with MSH2-MSH3 deficiency, might be near impossible given the vast majority of number of point mutations and small deletions associated with MMR deficiency are linked to polymerase slippage. Collectively, we provide a mechanistic explanation for how the MSH2-MSH3 complex facilitates efficient DSB repair by promoting HR via the recruitment of EXO1 and via preventing error prone TMEJ by blocking POLQ access.

## STAR METHODS

### KEY RESOURCES TABLE

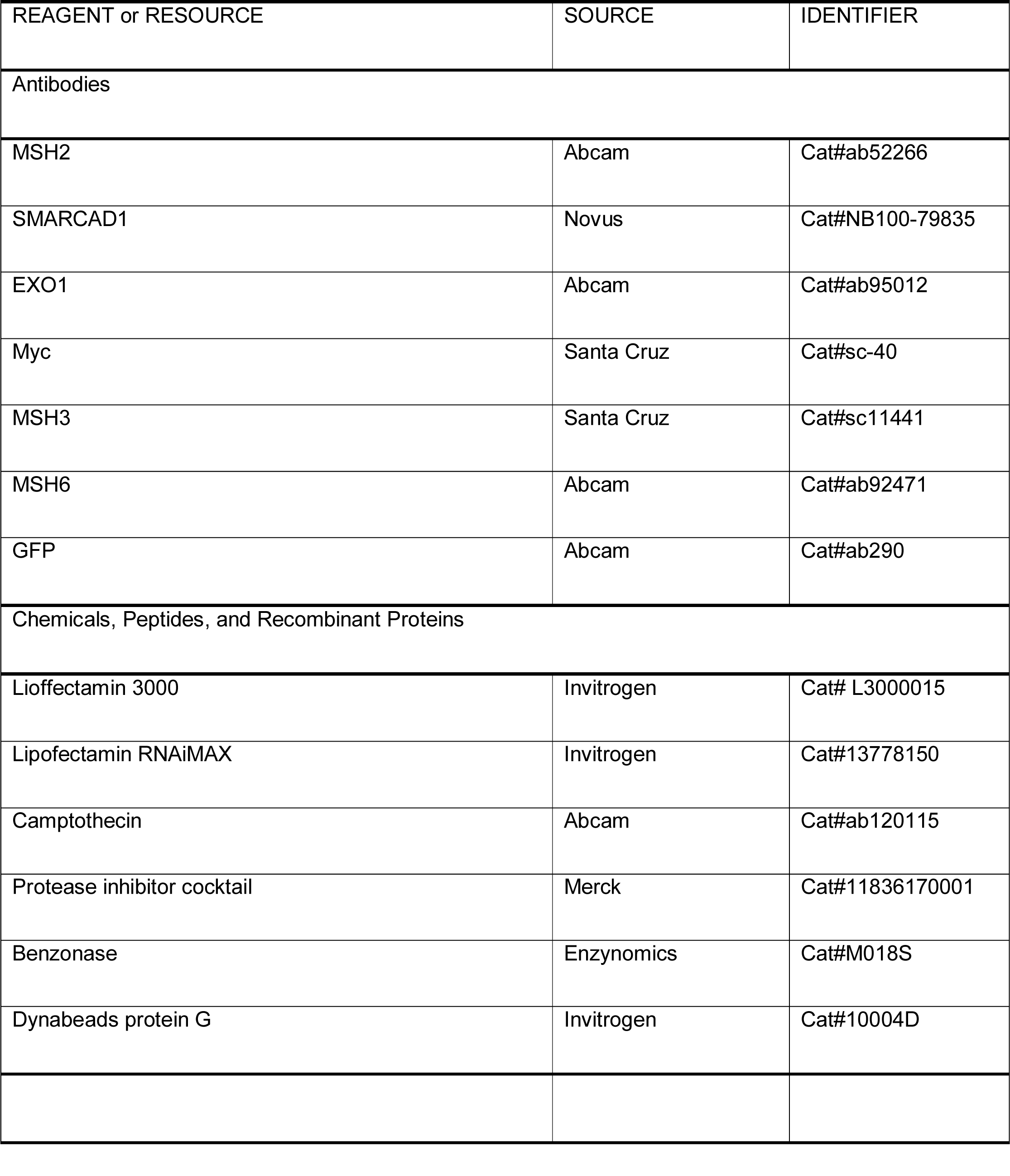
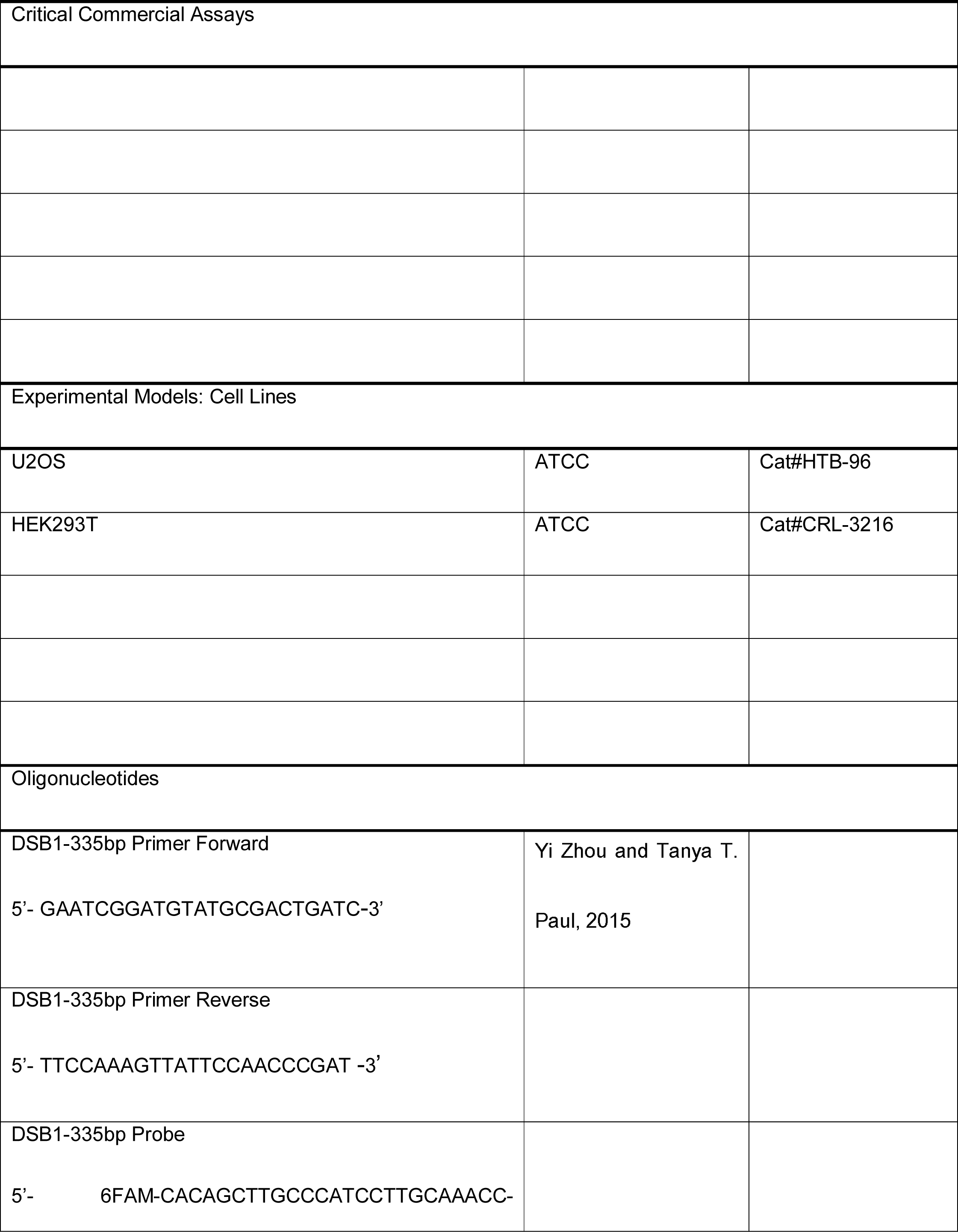
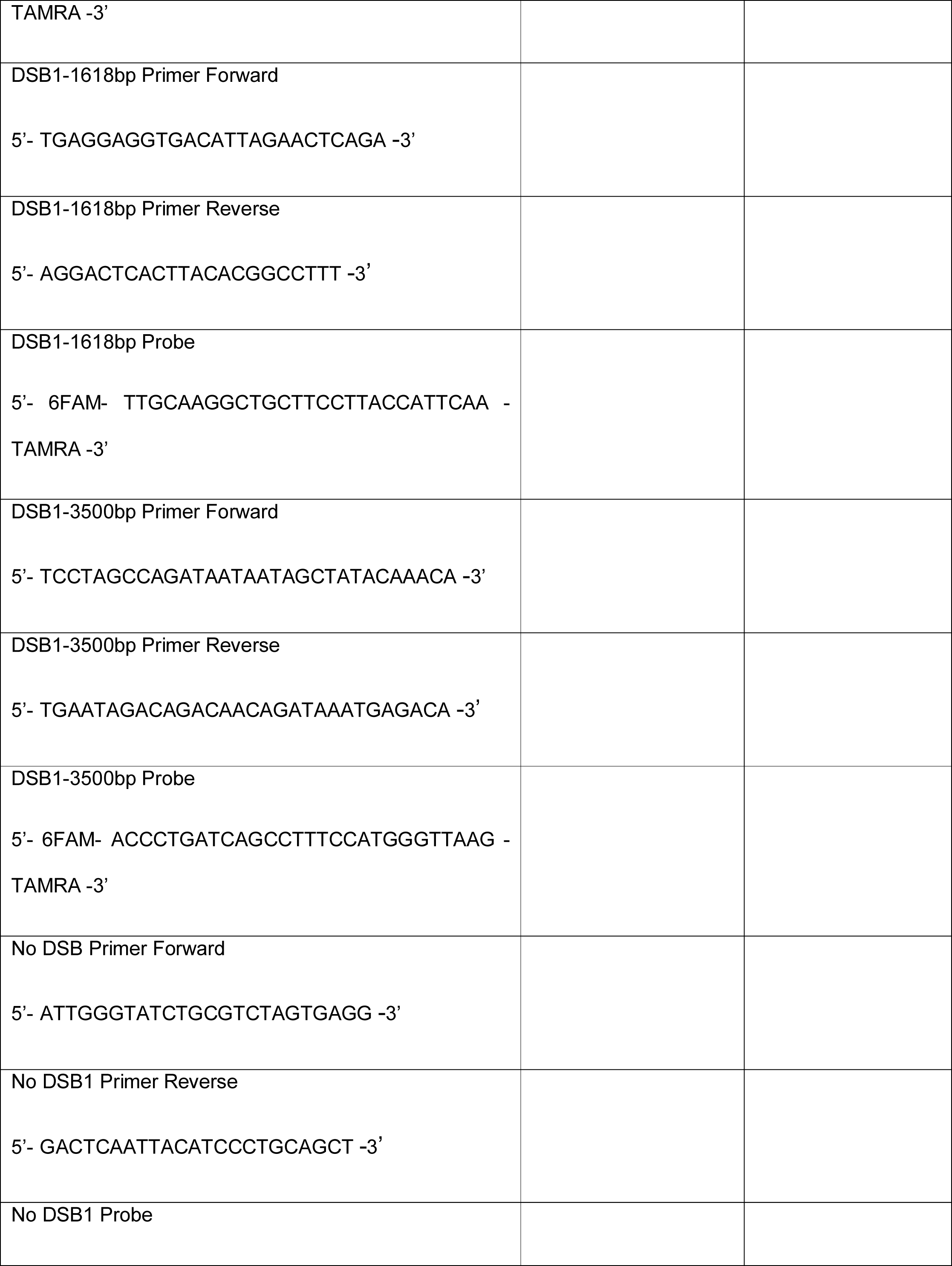
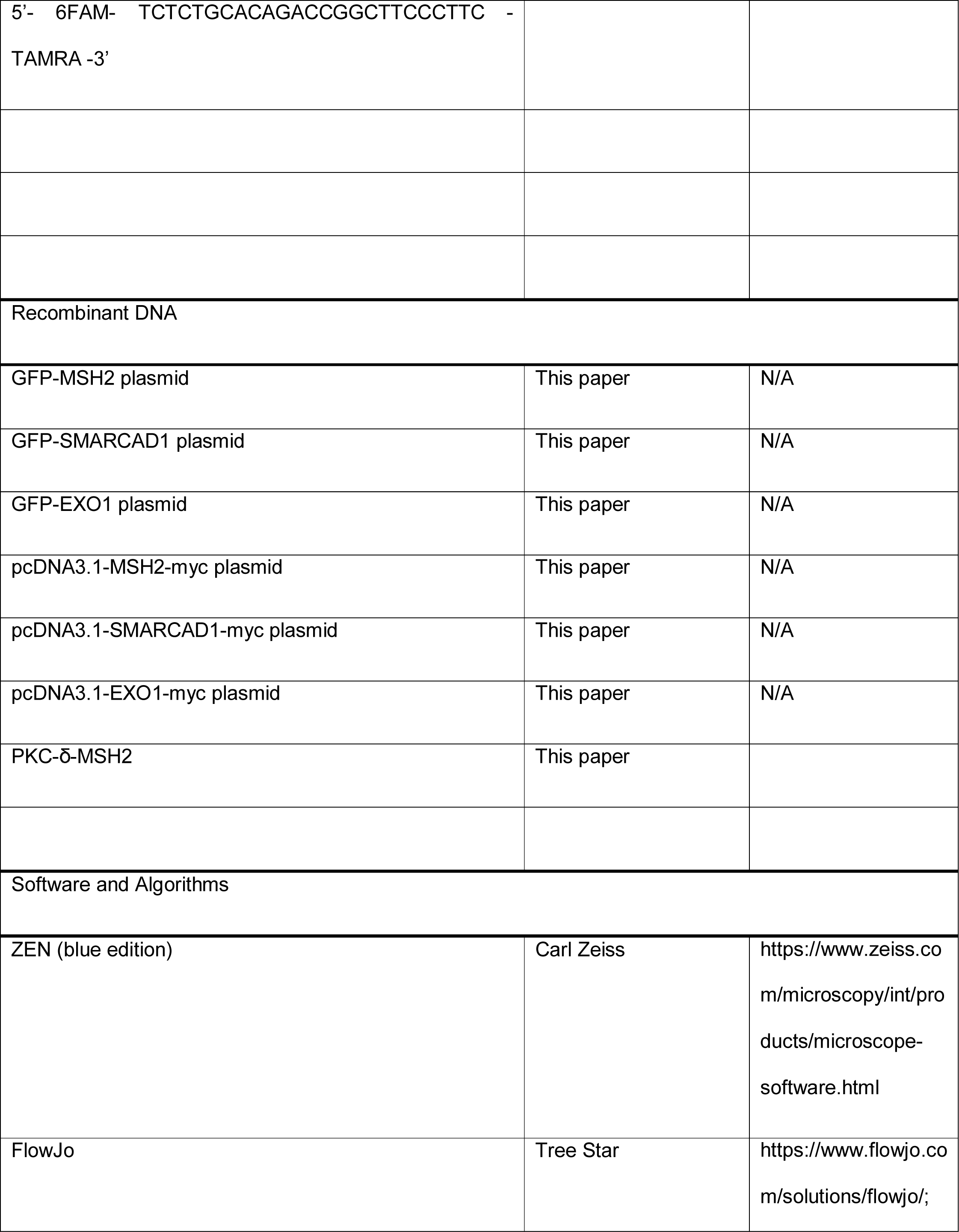
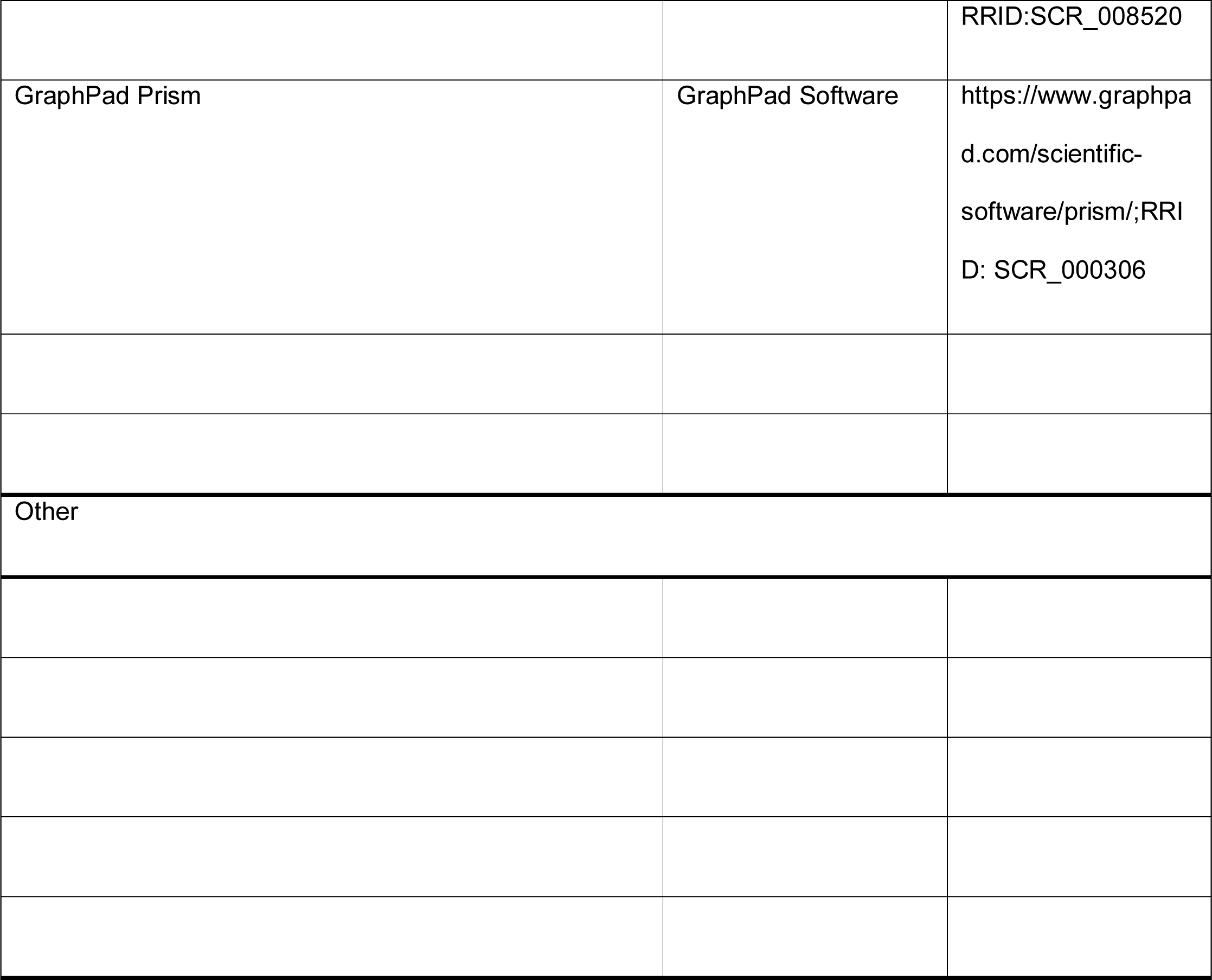

#### Lead contact and materials availability

Further information and requests for resources and reagents should be directed to and will be fulfilled by the lead Contact, yungjae Myung.

#### Cell culture and treatment

U2OS and HEK293T cells were purchased from ATCC and maintained in DMEM/ High Glucose (Hyclone) with 10% FBS (Millipore) and 1% penicillin and streptomycin (Invitrogen) at 37°C and 5% CO_2_. For DNA repair assays, U2OS cells stably expressing DR-℃GFP (HR), SA-GFP (SSA), EJ2-GFP (TMEJ) and EJ5-GFP (NHEJ) (Bennardo et al., 2008; Bennardo et al., 2009; Motegi et al., 2008; Pierce et al., 1999) were grown in DMEM (Gibco) containing 10% FBS (Merk) and 2μg/ml Puromycin (Invitrogen).

Human *MSH2* cDNA was PCR amplified from human cDNA isolated from HeLa cells using Trizol (Invitrogen) and cloned into EGFP-C2 vector using SalI and BamHI restriction sites and pcDNA3.1 myc-His A vector using BamHI and ApaI sites. SMARCAD1 cDNA [a gift from Tej K.Pandita (Chakraborty et al., 2018)] was cloned into EGFP-C2 vector using SalI and BamHI restriction sites and pcDNA3.1 myc-His A vector using BamHI and XbaI sites. EXO1 cDNA [a gift from Zhongsheng You (Chen et al., 2013)] was cloned into EGFP-C2 vector using SalI and BamHI restriction sites and pcDNA3.1 myc-His A vector using BamHI and ApaI sites. All cDNA were confirmed by sequencing.

The plasmid to express an active DNA polymerase fragment of POLQ (Sumo3 POLQM1) was a gift from Sylvie Doublie & Susan Wallace (Addgene plasmid # 78462) (Hogg et al., 2011). Full length POLQ without stop codon was cloned into plasmid pcDNA-DEST47 (Invitrogen) resulting in a protein tagged with GFP at the C terminus.

#### 355 nm laser microirradiation

We plated 3 × 10^5^ of U2OS cells in confocal dishes (SPL) and incubated 1 day. Then we transfected 2 μg of each plasmid expressing GFP-MSH2, GFP-SMARCAD1, or GFP-EXO1, with Lipofectamine 3000 according to the manufacturer’s instructions. Media containing plasmids with Lipofectamine were changed with 10 μM of 5-bromo-2’-deoxyuridine containing media after 4 h and incubated for 24 h. We used a 355 nm ultraviolet A laser for laser microirradiation and incubated cells in a 37°C chamber with 5% CO_2_. After each laser microirradiation, we obtained a cell image every 10 s during a 5 min interval with LSM880 confocal microscopy (Zeiss). The intensity of each laser strip was determined by Zen blue software (Zeiss) and the values were normalized with baseline values. At least 10 cells were used for quantification.

#### siRNA transfection

We transfected each 20 nM siRNA aliquot into cells using Lipofectamine RNAiMAX reagent (Invitrogen) and incubated for 48 h. The siControl (5’-CGU ACG CGG AAU ACU UCG A-3’), siMSH2 (5’-AAUCUGCAGAGUGUUGUGC-3’), siSMARCAD1 (5’-GACGAUUGAAGAAUCCAUGCU-3’), siEXO1 (5’-CAAGCCUAUUCUCGUAU-3’), and siMLH1 (5’-GUGUUCUUCUUUCUCUGUA-3’) oligonucleotides were purchased from Bioneer.

#### Plasmid transfection and immunoprecipitation

HEK293T cells were seeded to ∼60% confluence in 100-mm dishes and incubated 1 day. Each plasmid was then transfected with Transporter 5 (Polysciences) reagent according to the manufacturer’s instructions. After 24 h incubation, cells were washed with ice-cold phosphate-buffered saline (PBS) and lysed in buffer X (100 mM Tris-HCl pH8.0, 250 mM NaCl, 1 mM EDTA, 1% NP-40) with protease inhibitor cocktail (Merck, 11836170001), Benzonase (Enzynomics, M018S), and 5 mM MgCl2. We homogenized cell lysates by sonication and removed insoluble debris by centrifugation at 15000 rpm at 4°C for 10 min. Primary antibody (anti-Myc, 9E10, Santa Cruz) was added to the supernatant for overnight immunoprecipitation. The immunocomplexes were pulled down with dynabeads protein G beads (Invitrogen) and washed three times with buffer X. Samples were eluted with 2X NuPAGE sample buffer (Invitrogen), then subjected to SDS-PAGE. For endogenous immunoprecipitation, we used ∼80% confluent HEK293T cells and immunoprecipitated with each antibody (anti-MSH2, ab52266, abcam; anti-SMARCAD1, NB100-79835, Novus; anti-EXO1, ab95012, abcam).

#### *FokI* assay

We plated FokI-U2OS cells, a stable cell line with a *Fok*I restriction enzyme site, in four well plate and transfected with control or MSH2 siRNA. The next day, we used Lipofectamine 3000 to transfect with LacI-mCherry-FokI expression plasmid and GFP-*EXO1* or mNeon-*MRE11*. After 48 h, transfected cells were stained with Hoechst for 15 min to visualize the nucleus. Live cell images were obtained with LSM880 confocal microscopy (Zeiss).

#### Cell cycle analysis

We transfected U2OS cells in 60-mm diameter plate with the indicated siRNAs. After 48 h, we fixed cells with 70% (v/v) ice-cold ethanol and incubated at -20°C for 1 h. Then we washed cells with ice-cold PBS once and stained with propidium iodide in FACS buffer (1× PBS, 0.1% Triton X-100, 0.2 mg/ml RNase A) at 37°C for 30 min. Stained cells were analyzed using BD flow cytometry.

#### CUPID assay

We transfected HEK293T cells with *PKC-δ-MSH2* and either GFP-*SMARCAD1* WT or GFP-^SMARCAD1^-D1 mutant plasmids. For the MSH2 and EXO1 binding experiment, we transfected HEK293T cells with PKC-δ-*MSH2* and either GFP-*EXO1* WT or GFP-*EXO1* D13 mutant. After 24 h, cells were treated with 1 μM phorbol 12-myristate 13-acetate (PMA) and incubated for 5 min. We washed PMA treated cells two times with PBS, then incubated with 4% formaldehyde for 5 min and washed with PBS. We obtained cell images with LSM880 confocal microscopy.

#### HR, SSA, NHEJ, and MMEJ assays

SceI (pCAGGS-I-SceI, called pCBASce), empty vector (pCAGGS-BSKX), and dsRed vector (gift from Jeremy Stark) were prepared as described previously (Bennardo et al., 2008; Bennardo et al., 2009; Pierce et al., 1999). U2OS cells stably expressing DR-GFP, SA-GFP, EJ2-GFP, and EJ5-GFP plasmid were plated on a 12-well plate at 1×10^5^ cells/well. The following day, we transfected cells with 20 nM siRNA duplex mixed with RNAiMAX (Invitrogen) in Opti-mem. After 24 h, we performed the second round of transfection. To measure transient transfection efficiency after 2 days, we plated transfected cells on a 12-well plate at 1×10^5^ cells/well. The following day, we co-transfected cells with 0.5 µg of either I-SceI expression vector or empty vector, and 0.1 µg of dsRED vector (used as a transfection control) in 0.1 ml Opti-mem containing 3 µl of Lipofectamin 3000 (Invitrogen). After 6 h, we removed the media and replaced with growth media. Two days after I-SceI transfection, we analyzed the percentage of GFP+ cells using a Becton Dickinson FACSVerse flow cytometer. We calculated DNA repair efficiencies as previously described (Motegi et al., 2008). Experiments were repeated at least three times.

#### End resection assay

We trypsinized and centrifuged ER-*Asi*SI U2OS cells, then resuspended with 37°C pre-heated 0.6% low-melting agarose (Bio-Rad) in PBS (Sigma) at a concentration of 1.2 × 10^7^ cells/ml. We dropped 50 µl of cell suspension on a piece of Parafilm (Pechiney) to create a solidified agar ball, which we then transferred to a 2-ml Eppendorf tube. The agar ball was treated with 1 ml of ESP buffer (0.5 M EDTA, 2% N-lauroylsarcosine, 1 mg/ml proteinase K, 1 mM CaCl_2_, pH8.0) for 20 h at 16°C with rotation, followed by treatment with 1 ml of HS buffer (1.85 M NaCl, 0.15 M KCl, 5 mM MgCl_2_, 2 mM EDTA, 4 mM Tris, 0.5% Triton X-100, pH7.5) for 20 h at 16°C with rotation. After washing with 1 ml PBS (Sigma) for 6 h at 4°C with rotation, we melted the agar ball by placing the tube in a 68°C heat block for 10 min. The melted sample was diluted 15-fold with 68°C pre-heated ddH_2_O, mixed with an equal volume of the two designated NEB restriction enzyme buffers and stored at 4 C for future ° use. We measured the level of resection adjacent to specific DSBs by quantitative polymerase chain reaction (qPCR). The sequences of qPCR primers and probes are shown in the Key Resources table. We digested or mock digested 36 µl of genomic DNA sample with 80 units of restriction enzymes (BsrGI, HindIII-HF; New England Biolabs) at 37°C overnight. Three ° microliters of digested or mock digested samples were used as templates in 25 μl of qPCR reaction containing 12.5 μl of Taqman Universal PCR Master Mix (ABI), 0.5 mM of each primer, and 0.2 mM probe using a QuantStudio 7 Flex Real-Time PCR System (ABI). We determined the percentage of ssDNA (ssDNA%) generated by end resection at selected sites as previously described (Zhou and Paull, 2015). Briefly, for each sample, we calculated a Ct by subtracting the Ct value of the mock digested sample from the Ct value of the digested sample. We calculated ssDNA% with the following equation: ssDNA% = 1/(2^(4Ct-1)+0.5)*100 (Zhou and Paull, 2015).

#### RPA retention assay

Cells were treated 5 μ M of baicalein for 24 h. We transferred trypsinized cells to a 1.5 ml tube, washed with PBS, and permeabilized with 100 μl of 0.2% Triton X-100 in PBS for 10 min on ice. After washing with 1x PBS containing 1mg/ml of bovine serum albumin (PBS-BSA), and then cells were fixed and permeabilized with 100 µl of BD Cytofix/Cytoperm buffer (BD Biosciences) for 15min at room temperature. Fixed cells were washed with 0.5 ml of 1x BD Perm/Wash buffer (BD Biosciences) and suspended in 50 µl of 1x BD Perm/Wash buffer and incubated with anti-RPA2 and Alexa Fluor 488-secondary antibodies sequentially. The nucleus was visualized by PI staining for 15 min. Cells were analyzed using the Becton Dickinson FACSVerse flow cytometer.

#### DNA preparation for in vitro experiments

All DNA oligomers were chemically synthesized (Bioneer, South Korea) and listed in the Supplementary Table 1. Each set of oligomers were annealed by heating at 95°C for 20 min followed by slow cooling to 23°C.

#### Electrophoretic mobility shift assay (EMSA)

The EMSA assay for MSH2-MSH3 was performed by following the previous literature (Kumar et al., 2014). All reactions were performed at 23°C. 1 nM of Cy5-labeled 40 bp homoduplex, +8-loop DNA, or 58 bp flap DNA was incubated with MSH2-MSH3 at different concentrations in buffer H (20 mM HEPES [pH7.5], 100 mM NaCl, 1 mM DTT, 1 mM ATP, 2 mM MgCl_2_, 0.04 mg/ml BSA) for 5 min. For SMARCAD1, 1 nM of Cy5-labeled 40 bp +8-loop DNA or 58 bp flap DNA was incubated with SMARCAD1 at different concentrations in buffer H supplemented with 1 mM ATP for 15 min.

To test MSH2-MSH3 recruitment by SMARCAD1, 1 nM of Cy5-labeled 40 bp +8-loop DNA, or 58 bp flap DNA were incubated with 4 μM SMARCAD1 in buffer H supplemented with 1 mM ATP for 15 min. Then MSH2-MSH3 was added at different concentrations and incubated for 5 min. For EMSA assay for EXO1, 1 nM Cy5-labeled 40 bp homoduplex for DNA with +8-loop DNA was reacted with EXO1 at different concentrations (50, 100, 200, 400, 800, and 1600 nM) in buffer H for 20 min. To test EXO1 binding to MSH2-MSH3, 1 nM of Cy5-labeled 40 bp +8-loop DNA and 300 nM MSH2-MSH3 were incubated in buffer H for 5 min and EXOI was then added at different concentrations and further incubated for 20 min. All the reactants were then analyzed by running 5% non-denaturing PAGE at 130 V for 45 min in TE buffer (45 mM Tris-HCl [pH8.5], 0.5 mM EDTA) at 4°C. The gel was imaged by scanning Cy5 fluorescence using Typhoon RGB (GE Healthcare).

#### EXOI nuclease activity assay

The EXOI nuclease activity was tested with a previously established protocol (Daley et al., 2020). 20 nM of 40 bp DNA with 4-nt 3’ overhang labeled by Cy5 (40 bp overhang) was mixed with EXOI in EXOI buffer (20 mM HEPES [pH7.5], 100 mM KCl, 1 mM DTT, 100 μg/ml BSA, 0.05% Triton-X 100, 2 mM MgCl2) at different concentrations and incubated for 2 hours at 37°C. For deproteinization, SDS and proteinase K were added up to 0.2% and 0.25 μg/μl, respectively, and then further incubated for 20 min at 50°C. Digested DNA fragments were analyzed on 15% non-denaturing polyacrylamide gels in 0.5x TBE buffer at 200 V for 1 hour at 23°C. The gel was imaged by scanning Cy5 fluorescence using Typhoon RGB (GE Healthcare).

For the enhancement of EXO1 nuclease activity by MSH2-MSH3, 20 nM of 90 bp flap DNA labeled by Cy5, which had a flap of dT_18_ and a 15-nt gap, was mixed with 500 nM MSH2-MSH3 and incubated for 5 min at 23°C. Then EXOI was added to the MSH2-MSH3-flap DNA complex in reaction buffer (20 mM HEPES [pH7.5], 100 mM NaCl, 1 mM DTT, 2 mM MgCl_2_, 1 mM ATP, and 40 μg/ml BSA) at different concentrations (0.25, 0.5, 1, 2, 4, 8 μM) and incubated for 2 hours at 37°C. The reaction was stopped and deproteinized with 0.2% SDS and 0.25 μg/μl proteinase K and then further incubated for 20 min at 50°C. Digested DNA fragments were analyzed in 8% non-denaturing polyacrylamide gels in 0.5X TBE buffer at 150 V for 70 min at 4°C. The gels were imaged using Typhoon RGB (GE Healthcare).

#### Protein purification

Full length human MSH2-MSH3, EXO1 and SMARCAD1 were expressed by infecting Hi5 insect cells with each amplified baculoviruses. After 48 hours of virus infection, cells were harvested. Cells were resuspended in buffer A containing 25 mM sodium phosphate, pH 7.8, 400 mM NaCl, 10 mM imidazole (40 ml per 1l cell). Supernatant were applied on a Ni^2+^ affinity HiTrap chelating HP column (GE Healthcare Life Sciences, 17040901). Proteins were eluted with linear gradient of buffer B (buffer A + 400 mM imidazole). Protein peaks were collected and concentrated using Amicon ultra-15 50K centrifugal filter. Concentrated proteins were further applied to a HiLoad 16/600 Superdex 200 pg column (GE Healthcare Life Sciences, 28989335) equilibrated in buffer consisting of 25 mM Tris-HCl, pH 7.5, 150 mM NaCl, 5 mM dithiothreitol (DTT). Fractionated protein peak of each step was confirmed by SDS-PAGE and concentrations of proteins were measured by Bradford assay.

#### DNA polymerase assays

An active DNA polymerase fragment of POLQ was expressed from the Sumo3 POLQM1 plasmid and purified as described (Hogg et al., 2011). Klenow Fragment (3′→5′ exo-) was purchased from NEB. POLQ was diluted in buffer containing 37.5 mM Tris-HCl, pH8.0, 40 mM NaCl, 2.5 mM DTT, 6.25% glycerol, 0.0125% Triton X-100, and 0.125% bovine serum albumin (BSA). Klenow Fragment (3′→5′ exo-) was diluted in buffer containing 50 mM Tris-HCl pH7.5, 1 mM DTT, 50% Glycerol and 0.1 mM EDTA. PAGE-purified oligonucleotides were purchased from IDT. Primer (5’-AAAAAAAAATATGATG) was 5′-labeled using polynucleotide kinase and [γ-^32^P]dATP. 5′-^32^P-labeled primer was annealed to template oligonucleotides containing no or 2 bp mismatch bases in different positions as follows (2 bp mismatch underlined): No-mismatch template: TTTTTTTTTATACTACTACTACGACTGCTC-5. MM-1,2 template: TTTTTTTTTATACTGTTACTACGACTGCTC-5′. MM-3,4 template: TTTTTTTTTATATCACTACTACGACTGCTC-5′. MM-5,6 template: TTTTTTTTTACGCTACTACTACGACTGCTC-5′. POLQ reaction mixtures (10 μl) containing 30 mM Tris-HCl pH8.0, 0.5% glycerol, 0.4 mM dithiothreitol (DTT), 0.02% BSA, 20 mM MgCl2, 100 μM of each dNTP, 100 nM of the primer-template or primer. Klenow Fragment (3′→5′ exo-) reaction mixtures (10 μl) contained 50 mM Tris-HCl pH7.2, 0.1 mM DTT, 10 mM MgSO4, 100 μM of each dNTP, and 100 nM of the primer-template. After incubation at 37°C for 10 min, reactions were terminated by adding 10 μl of formamide stop buffer (98% formamide, 10 mM EDTA pH8.0, 0.025% xylene cyanol FF, 0.025% bromophenol blue) and boiling at 95°C for 3 min. Products were electrophoresed on a denaturing 20% polyacrylamide (7 M urea gel) and analyzed with Typhoon RGB (Amersham). For the assay with MutSβ derivatives, MutSβ derivatives were incubated first with DNA substrates in the reaction buffer without POLQ at room temperature (25°C) for 20 min and incubated with POLQ at 37°C for 10 min.

#### Determination of termination probability and amount of full-length extension

The termination probability at position N was defined as the band intensity at N divided by the total intensity for all bands ≥N, as previously described (Kokoska et al., 2003). Quantification of full-length extension products was defined as the fully extended band intensity divided by the intensity for all bands ≥ N0 (Primer position).

#### Targeted deep sequencing

We transfected 20 nM of control or MSH2 siRNA to 1.5 × 10^6^ HEK293T cells in 10 cm dish. After 24 h incubation, we transfected 4 μg of p3s-Cas9-HN and 6 μg of each plasmid expressing mCherry-gRNA (targeting CEL and non-targeting gRNA control 1 and control 2) with lipofectamine 3000 according to the manufacturer’s instructions. Cells were incubated for 2 days and only Cas9 and mCherry-gRNA transfected cells were sorted by FACS using FACSAria Fusion (BD bioscience). 2 × 10^5^ cells expressing mCherry signal were sorted and genomic DNA were extracted by QiAamp DNA Mini Kit (Qiagen) according to the manufacturer’s instructions. To measure the mutation frequency at the CRISPR-Cas9 induced DSB site by genomic sequencing, we started nested PCR with 200 ng of genomic DNA using each primer, CEL F1: 5’-TGTGGACATCTTCAAGGGCA-3’, CEL R1: 5’-AGATCATAACGGGCAGGTCC-3’, CEL F2: 5’-GTGACTGGAGTTCAGACGTGTGCTCTTCCGATCT CCTTCCTCATGCCAACTCCT-3’, CEL R2: 5’-ACACTCTTTCCCTACACGACGCTCTTCCGATCTCTCAAGCCAGGAGTAGACCC, Pooled PCR products were sequenced using NextSeq 500/550 Mid Output kit v2.5, 300 cycles (Illumina). Sequencing reads were analyzed using CRISPRpic (Lee et al., 2020) software for frequency of microhomology mediated repair.

## Supporting information

all six supplementary figures

## Acknowledgments

We thank the members of the IBS Center for Genomic Integrity, especially Drs. Anton Gartner and Kyoo-young Lee for their helpful comments and discussion. We thank Drs. Sylvie Doublie and Susan Wallace for providing the plasmid, Sumo3 POLQM1 (Addgene plasmid # 78462). We thank Dr. Jeremy Stark for providing cells to measure HR, SSA, NHEJ, and MMEJ. This research was supported by the Samsung Science and Technology Foundation under Project Number SSTF-BA1901-13 to JY Lee, the National Research Foundation (grant number NRF-2021R1C1C1005358 and NRF-2018R1A5A2023879) to JM Oh and the Institute for Basic Science (grant number IBS-R022-D1) to K Myung.

## Notes

### Competing Interest Statement

The authors have declared no competing interest.

