## Supplementary figures and images for "MSH2-MSH3 promotes DNA end resection during HR and blocks TMEJ through interaction with SMARCAD1 and EXO1"

### all six supplementary figures

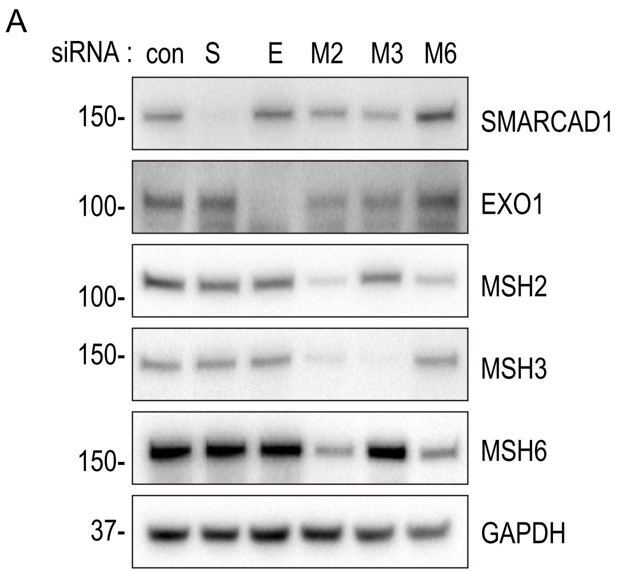


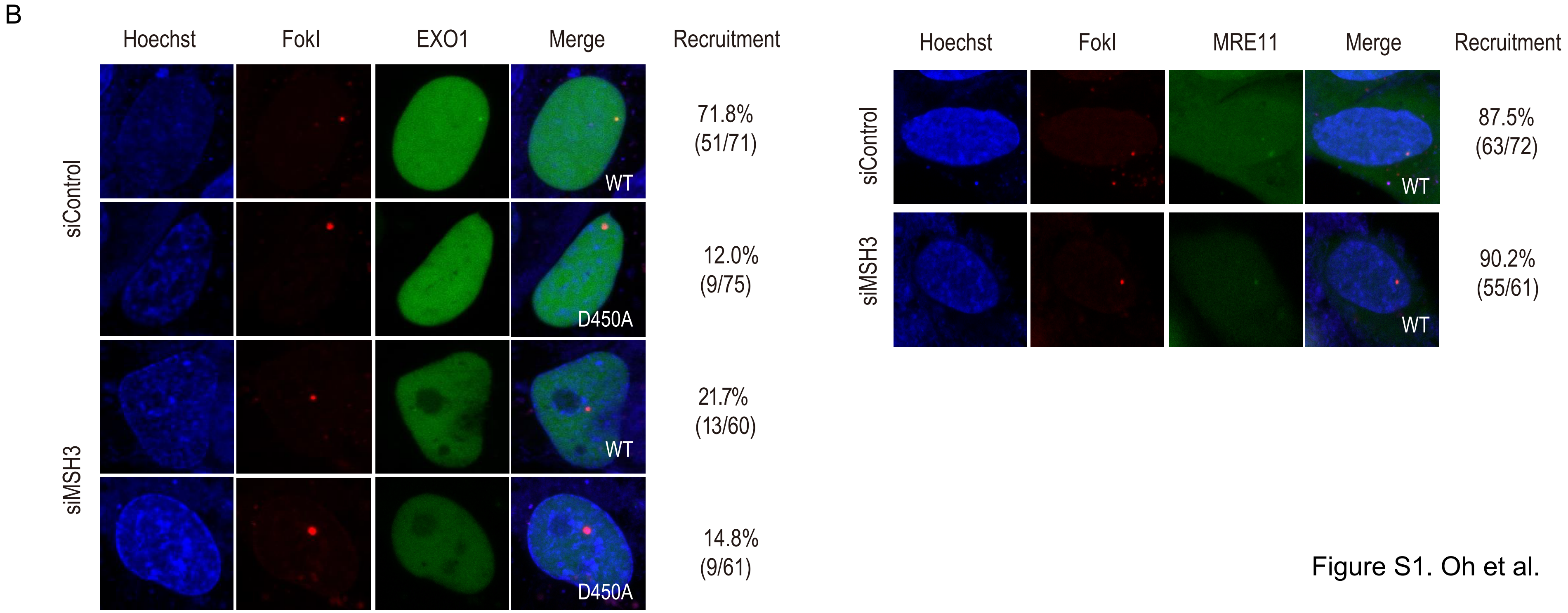


Figure S1. Oh et al.

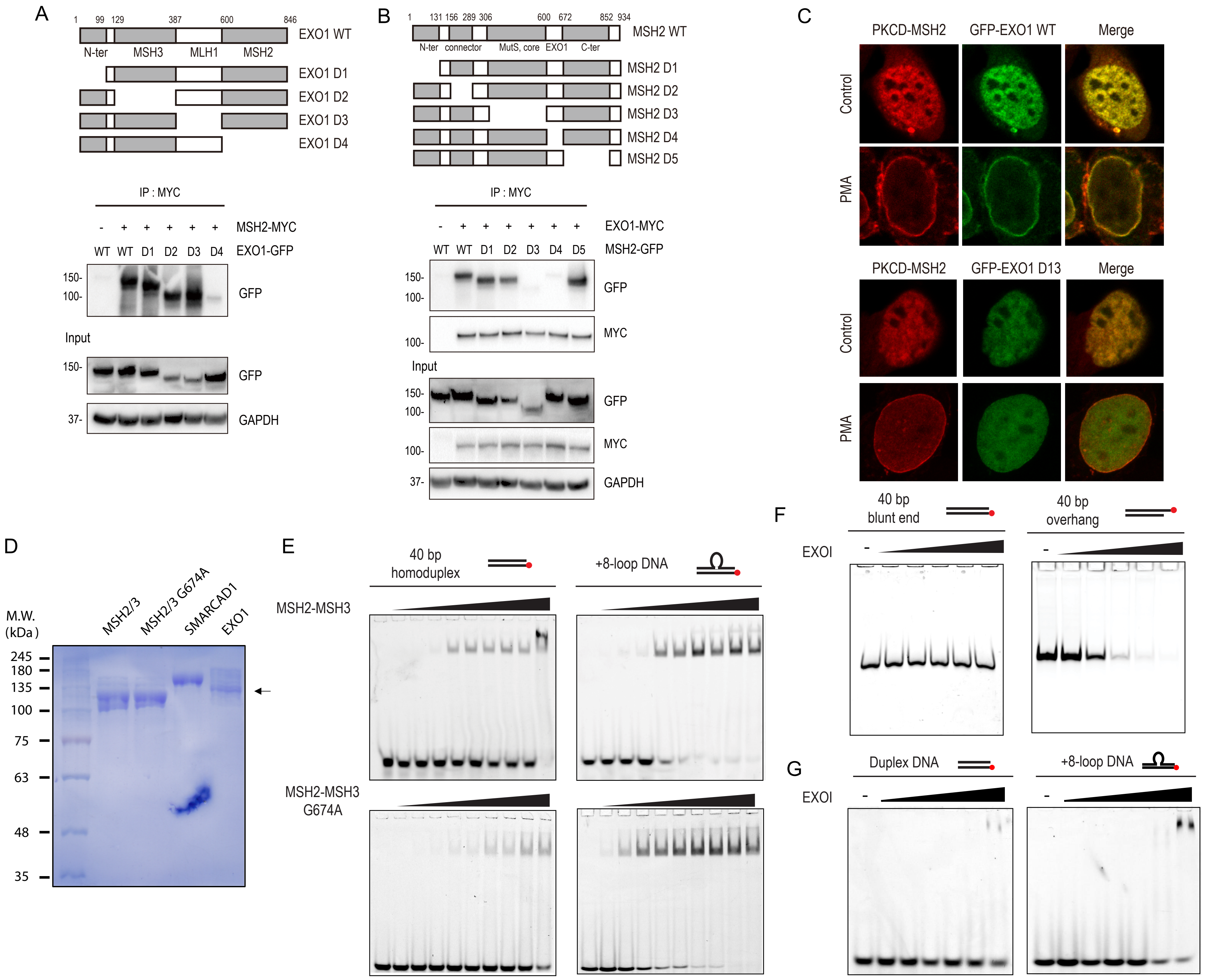


Figure S2. Oh et al.

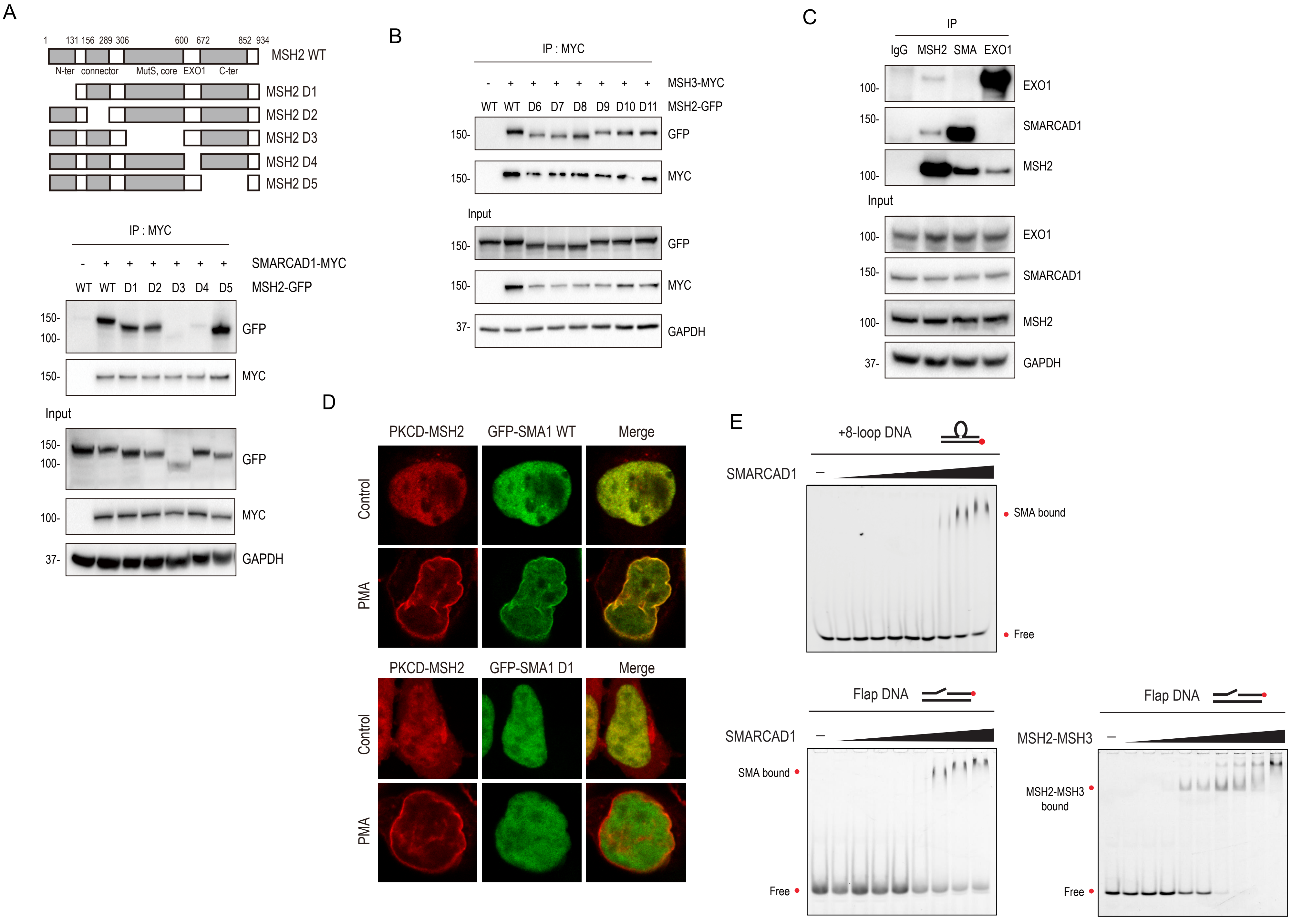


Figure S3. Oh et al.

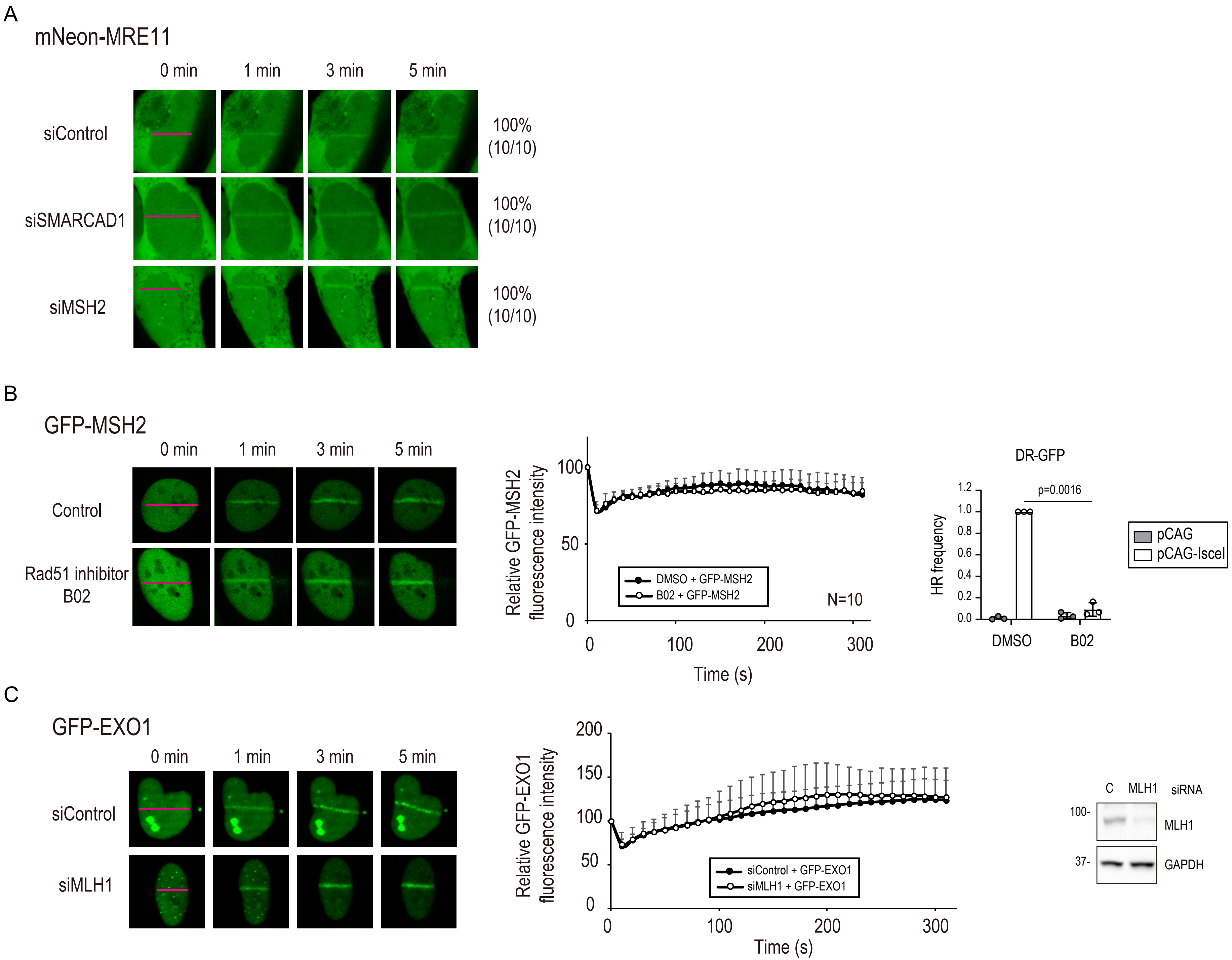


Figure S4. Oh et al.

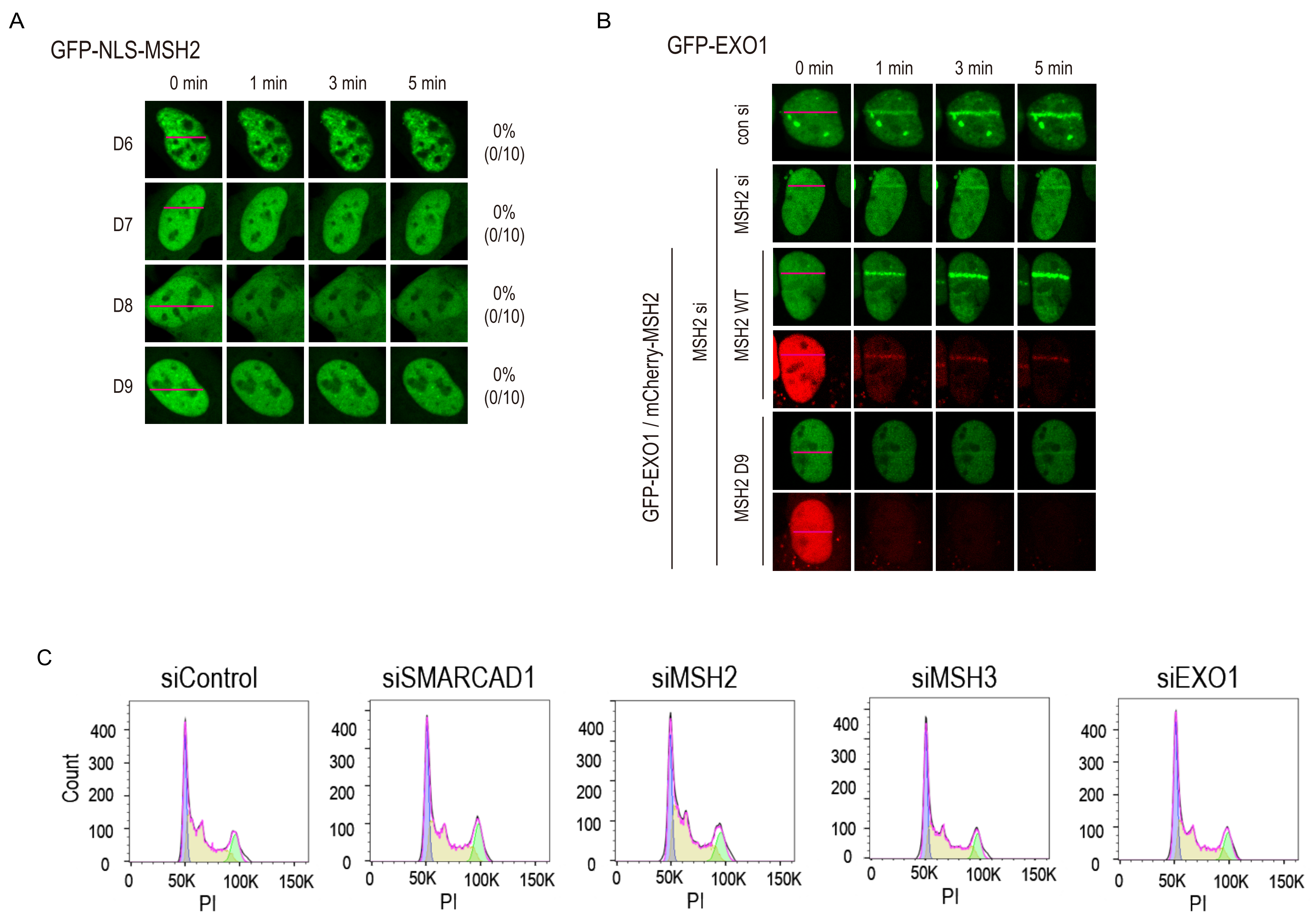


Figure S5. Oh et al.

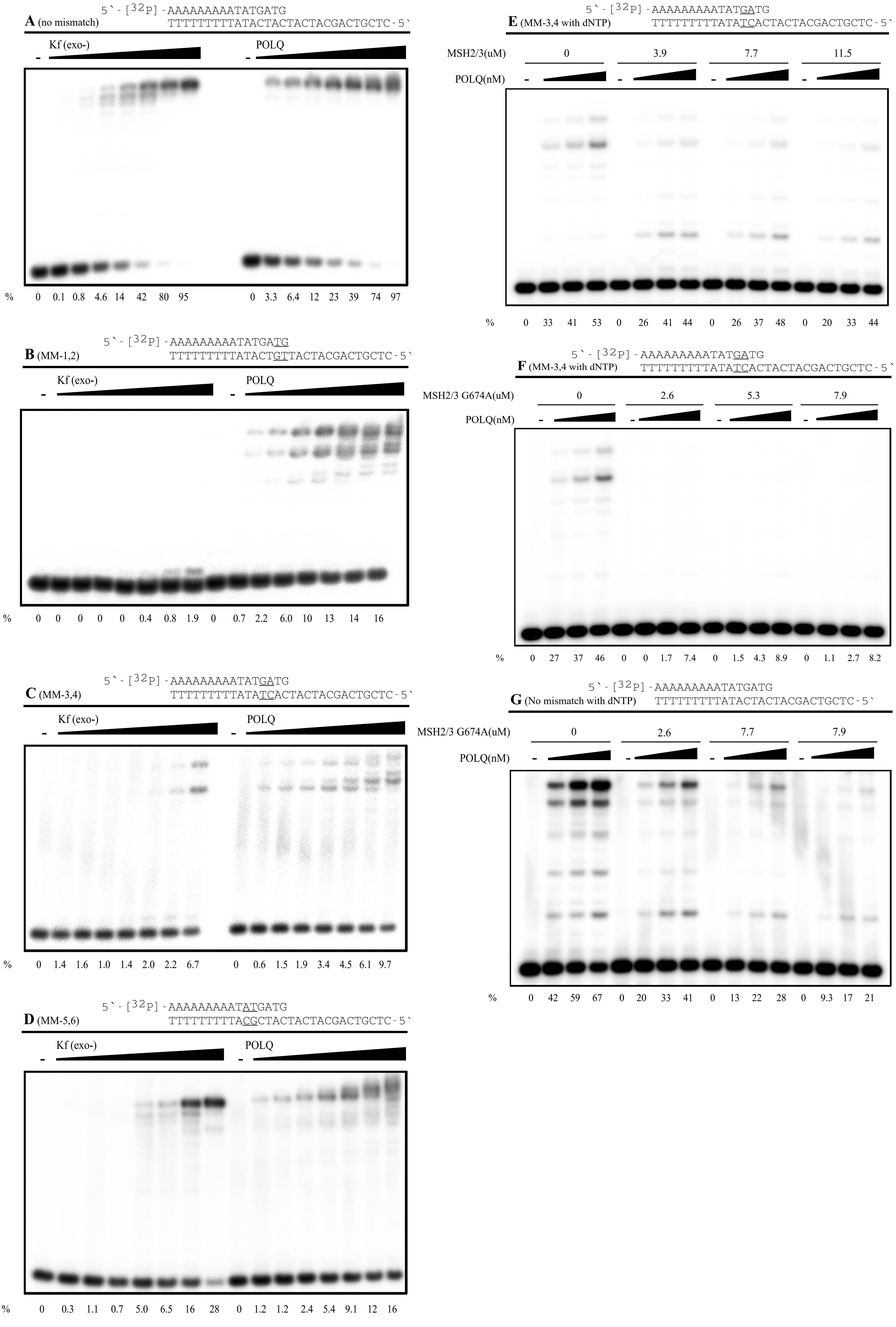
