## supplementary figure legends and table for "MSH2-MSH3 promotes DNA end resection during HR and blocks TMEJ through interaction with SMARCAD1 and EXO1"

Supplementary Information

Jung-Min Oh^1,2^*, Yujin Kang^3^*, Jumi Park^3^, Yubin Sung^1^, Dayoung Kim^4^, Yuri Seo^1,5^, Eun A Lee^1^, Jae Sun Ra^1^, Enkhzul Amarsanaa^1,3^, Young-Un Park^1,6^, Hongtae Kim^1,3^, Orlando Schärer^1,3^, Seung Woo Cho^1,4^, Changwook Lee^3^, Kei-ichi Takata^1,3^, Ja Yil Lee^1,3+^ and Kyungjae Myung^1,4+^

1. Center for Genomic Integrity, Institute for Basic Science (IBS), Ulsan 44919, Republic of Republic of Korea.

2. Department of Oral Biochemistry, Dental and Life Science Institute, School of Dentistry, Pusan National University, Yangsan 50612, Republic of Korea

3. Department of Biological Sciences, Ulsan National Institute of Science and Technology, Ulsan 44919, Republic of Korea

4. Department of Biomedical Engineering, Ulsan National Institute of Science and Technology, Ulsan 44919, Republic of Korea

5. Current address; Chungnam National University, Daejeon, Republic of Korea

6. Current address; CasCure Therapeutics, Ulsan, Republic of Korea

*Equal contributions

+corresponding authors

 (Ja Yil Lee) or (Kyungjae Myung)

**Supplementary Figure Legends**

Supplementary figure 1. MSH2 Depletion does not affect MRE11 Recruitment

(A) U2OS cells were transfected with each siRNA and incubated for 48 hours. Cells were harvested and each protein expression was analyzed by Western blot. (B) GFP-EXO1 or mNeon-MRE11 recruitment to *Fok*I induced DSB sites was measured in control and MSH3 knocked down cells.

Supplementary Figure 2. EXO1 interacts with MSH2.

(A) Diagram of EXO1 WT and deletion mutants. Co-IP was performed after 24hours of co-transfection of myc-MSH2 WT and each GFP-EXO1 WT or deletion mutant to HEK293T cells. (B) Diagram of MSH2 WT and deletion mutants. HEK293T cells co-transfected with myc-EXO1 WT and each GFP-MSH2 WT or mutant were subjected to co-IP analysis. (C) HEK293T cells were co-transfected with PKC-δ-MSH2 and GFP-EXO1 WT or GFP-EXO D13. Cells were treated with 1 μM PMA and incubated for 5 min, then images were taken by confocal microscopy. (D) Coomassie Brilliant Blue stained 8% Tris-Glycine SDS-PAGE gels of purified proteins used in this study. Black arrow indicates EXO1. (E) DNA binding analysis for MSH2-MSH3 or MSH2 (G674A)-MSH3 with 40 bp homoduplex (left) and +8-loop DNA (right) at different concentrations (0, 10, 20, 40, 80, 100, 150, 200, 300, and 400 nM). Both 40 bp homoduplex and +8-loop DNA have Cy5 at 5’ end of one strand. MSH2-MSH3 shows higher binding affinity to +8-bubble than homoduplex. (F) Exonuclease activity analysis for EXO1 at different concentrations (0, 200, 400, 800, 1600, and 3200 nM). 40 bp DNA with 4-nt 3’-overhang labeled with Cy5 was used as DNA substrate. DNA substrate is degraded due to the digestion by EXO1 in dose dependent manner. (G) EMSA for EXO1. Duplex or +8-loop DNA was reacted with EXO1 at different concentrations (0, 50, 100, 200, 400, 800, and 1600 nM).

Supplementary figure 3. SMARCAD1 interacts with MSH2 not EXO1.

(A) Diagram showed MSH2 WT and deletion mutants. HEK293T cells were co-transfected with myc-SMARCAD1 WT and each GFP-MSH2 WT or mutants. (B) HEK293T cells were co-transfected with myc-MSH3 WT and GFP-MSH2 WT or mutant to check the interaction between MSH2 and MSH3. (C) HEK293T cells were harvested and immunoprecipitated with IgG, MSH2, SMARCAD1 or EXO1 antibody to check the interaction between MSH2, SMARCAD1 and EXO1. (D) HEK293T cells were co-transfected with PKC-δ-MSH2 and GFP-SMARCAD1 WT or GFP-SMARCAD1 D1. 1 μM PMA was treated for 5min before taking images under confocal microscopy. (E) DNA binding analysis for SMARCAD1 with +8-loop DNA at different concentrations (0, 50, 100, 200, 400, 800, 1000, 2000, 4000, and 8000 nM, top). DNA binding analysis for SMARCAD1 with 58 bp flap DNA at different concentrations (0, 400, 800, 1000, 2000, 4000, 8000, 16000 and 32000 nM, bottom left). DNA binding analysis for MSH2-MSH3 with 58 bp flap DNA at different concentrations (0, 10, 20, 40, 80, 100, 150, 200, 300, and 400 nM, bottom right).

Supplementary figure 4. MRE11 recruitment to DSB sites is not affected by SMARCAD1 and MSH2 knocked down.

(A) U2OS cells were transfected with control, SMARCAD1 or MSH2 siRNA. 24 hours later, mNeon-MRE11 was transfected then, cells were incubated with 10 μM of BrdU for 24 hours before monitor the MRE11 recruitment to microirradiation induced DSB sites. (B) U2OS cells were transfected with GFP-MSH2 and incubated with 10 μM of BrdU for 24 hours. RAD51 inhibitor B02 was incubated with last 4 hours before microirradiation. The effect of B02 was confirmed by HR assay. (C) MLH1 knock down, which was confirmed by western blot, did not affect GFP-EXO1 recruitment to microirradiation induced DSB sites.

Supplementary figure 5. EXO1 recruitment to DSB sites depends on MSH2

(A) NLS sequences were attached to each GFP-MSH2 mutant from D6 to D9. U2OS cells were transfected with each mutant and incubated with 10 μM of BrdU for 24 hours. (B) Control and MSH2 knocked down U2OS cells were co-transfected with GFP-EXO1 WT and either mCherry-MSH2 WT or mCherry-MSH2 D9 mutant to monitor their recruitment to the microirradiation induced DSB sites. (C) Indicated each siRNA was transfected to U2OS cells for 48 hours and cell cycle was measure by FACS analysis.

Supplementary Figure 6. POLQ efficiently extends from a mismatched primer end.

Increasing amounts of POLQ (0.3, 0.6, 1.3, 0.25, 0.5, 10, 20 and 40 nM), and KF (exo-) (3.9, 7.8, 15.6, 31.3, 62.5, 125, 250 and 500 fM) were incubated with the 5′-^32^P-labeled primer templates indicated above the gel in the presence of all 4 nt at 37°C for 10 min. The first lane contained no enzyme (-). The percentage (%) of the product extension from the primer is shown below each lane. (A) no mismatch. 2 mismatched base pairs were placed at 1 and 2 bp (B), 3 and 4 bp (C), and 5 and 6 bp (D) from the primer-template junction.

(E-G) Increasing amounts of POLQ (0.3, 0.6, 1.3 nM) were incubated in the presence of indicated amount of wild-type or mutant MSH2-MSH3 with the 5′-^32^P-labeled primer templates indicated above the gel in the presence of all 4 nt at 37°C for 10 min. The first lane contained no enzyme (-). The percentage (%) of the product extension from the primer is shown below each lane. 2 mismatched base pairs were placed at 3 and 4 bp from the primer-template junction. (E) POLQ was incubated with wild-type MSH2-MSH3 and mismatched substrate (E), mutant MSH2-MSH3 and mismatched substrate (F), or mutant MSH2-MSH3 and non-mismatched substrate (G).

**Supplementary Table. List of oligomers**

| Name | Sequence |
| --- | --- |
| 40 bp  homoduplex | 5’-ACCGAATTCTGACTTGCTAGGACATCTTTGCCCACGTTGA-3’ |
|  | 5’-**Cy5**-TCAACGTGGGCAAAGATGTCCTAGCAAGTCAGAATTCGGT-3’ |
| +8-loop DNA* | 5’-ACCGAATTCTGACTTGCTAGGTGTGTGTGACATCTTTGCCCACGTTGA-3’ |
|  | 5’-**Cy5**-TCAACGTGGGCAAAGATGTCCTAGCAAGTCAGAATTCGGT-3’ |
| 40 bp  overhang** | 5’-ACCGAATTCTGACTTGCTAGGACATCTTTGCCCACGTTGATTTT-**Cy5**-3’ |
|  | 5’-TCAACGTGGGCAAAGATGTCCTAGCAAGTCAGAATTCGGT-3’ |
| 58 bp flap DNA*** | 5’- GTGCACTCTCAGTACAATCTGCTCTGATGC**TTTTTTT** -3’ |
|  | 5’- TAAGCCAGCCCCGACACCCG **Cy5**-3’ |
|  | 5’- CGGGTGTCGGGGCTGGCTTAACTATGCGGCATCAGAGCAGATTGTACTGAGAGTGCAC -3’ |
| 90 bp flap DNA*** | 5’-ACGCATCTGTGCGGTATTTCACACC**TTTTTTTTTTTTTTTTTT**-**Cy3**-3’ |
|  | 5’-TCAGTACAATCTGCTCTGATGCCGCATAGTTAAGCCAGCCCCGACACCCG-**Cy5**-3’ |
|  | 5’-CGGGTGTCGGGGCTGGCTTAACTATGCGGCATCAGAGCAGATTGTACTGAGAGTGCACCATATGCGGTGTGAAATACC  GCACAGATGCGT-3’ |

* Red represents 8-nt bubble.

** Blue represents 4-nt single-stranded overhang.

*** Bold represents single-stranded flap.
